# Generation of lineage-resolved complete metagenome-assembled genomes by precision phasing

**DOI:** 10.1101/2021.05.04.442591

**Authors:** Derek M. Bickhart, Mikhail Kolmogorov, Elizabeth Tseng, Daniel M. Portik, Anton Korobeynikov, Ivan Tolstoganov, Gherman Uritskiy, Ivan Liachko, Shawn T. Sullivan, Sung Bong Shin, Alvah Zorea, Victòria Pascal Andreu, Kevin Panke-Buisse, Marnix H. Medema, Itzik Mizrahi, Pavel A. Pevzner, Timothy P.L. Smith

**Affiliations:** USDA Dairy Forage Research Center, Madison, WI 53593; University of California, San Diego, CA; Pacific Biosciences, Menlo Park, CA; St. Petersburg State University, St. Petersburg, Russia; Phase Genomics, Seattle, WA; USDA Meat Animal Research Center, Clay Center, NE; Ben Gurion University of the Negev, Beer Sheba, Israel; Bioinformatics Group, Wageningen University, Wageningen, the Netherlands

**Author notes:** These authors contributed equally to this manuscript (Derek M. Bickhart, Mikhail Kolmogorov).

## Abstract

Microbial communities in many environments include distinct lineages of closely related organisms which have proved challenging to separate in metagenomic assembly, preventing generation of complete metagenome-assembled genomes (MAGs). The advent of long and accurate HiFi reads presents a possible means to address this challenge by generating complete MAGs for nearly all sufficiently abundant bacterial genomes in a microbial community. We present a metagenomic HiFi assembly of a complex microbial community from sheep fecal material that resulted in 428 high-quality MAGs from a single sample, the highest resolution achieved with metagenomic deconvolution to date. We applied a computational approach to separate distinct haplotype lineages and identified haplotypes of hundreds of variants across hundreds of kilobases of genomic sequence. Analysis of these haplotypes revealed 220 lineage-resolved complete MAGs, including 44 in single circular contigs, and demonstrated improvement in overall assembly compared to error-prone long reads. We report the characterization of multiple, closely-related microbes within a sample with potential to improve precision in assigning mobile genetic elements to host genomes within complex microbial communities.

## Introduction

The creation of reference-quality, species-level assemblies from metagenome communities is exceedingly difficult. In particular, generating a complete genome assembly of a microbe closely related to a more abundant member of the community has been an elusive goal. Previous short-read studies have resulted in high-quality metagenome-assembled genomes (MAG)^1^ only after extensive polishing and manual curation of initial contigs^2^. However, if a community contains thousands of organisms at different levels of abundance, manual curation of each MAG to achieve reference quality is extremely laborious. Generally, the MAGs assembled from short reads are represented by hundreds or even thousands of contigs, many of which have fragmented open reading frames (ORFs) at their ends. A major source of discontinuity in metagenome assembly appears to be from the prevalence of high sequence identity orthologous genes and operons^3^. These genomic features tend to be repetitive in the community and can preclude complete assembly unless the data include sequencing reads that span the entire shared region. Furthermore, nearly all short-read and long-read assembly algorithms typically collapse the variant features into a single representation that does not reflect the true strain- or species-level diversity of a subpopulation within the community^4,5^, moreover, consensus assemblies might include various artifacts arising from the variation collapsing procedure, e.g. frame shifts, complicating downstream analysis^6^. Ambiguity resulting from the metagenomic assembly of short-reads or error-prone long-reads^7^ has therefore left the possibility of first-pass characterization of microbial strains out of reach.

Generation of high-quality assemblies of individual microbial lineages within metagenomes remains a substantial challenge. Binning methods were developed to address issues with assembly fragmentation and organize contigs into candidate MAGs based on assumptions of shared sequence composition^8^ or orthologous linkage data^9^. The presence of single copy genes (SCG) expected to be in all bacterial and archaeal lineages has been proposed as a measure of the completeness and redundancy within these bins^2^. High-quality draft MAGs are defined in the literature as having over 90% of the expected count of SCG with less than 5% redundancy of their prevalence^1^. However, bacterial and archaeal lineages may contain significant accessory gene content^10^ that is not assessed using these metrics. Even though bins are often generalized to represent distinct microbial taxonomic units in a sample, they are rarely assumed to accurately represent true, genetically distinct microbial populations in a sample. This problem has been addressed by multiple studies^11,12^, and precise definitions for individual, highly resolved MAGs remain contextual to each study. Similar to one of these studies^11^, we focus on generating separate representative reference genomes for distinct microbial lineages within an individual metagenome, which we define as “lineage-resolved MAGs”^1,13^. Combined with prior definitions of SCG quality metrics, we further extend the term to “lineage-resolved complete MAGs” for all such assemblies which have high degrees of SCG completeness (> 90%), low degrees of SCG redundancy (< 10%), and the separation of all observable variant lineages of microbial taxa into individual MAGs. Tools have been developed to identify or separate lineage-resolved complete MAGs from metagenomic bins post-hoc, but these tools often rely on co-assembly data, assembly graphs or various statistical methods to overcome biases in read-alignments to estimate strains from observed genetic variant data and therefore require more curation to properly disentangle lineages from MAGs^11,14,15^. Furthermore, these workflows are designed primarily to identify strain lineages from alignments of short-read data and do not capture variant linkage data from longer read datasets. A recent attempt to adapt uncorrected long reads to this purpose requires the use of manual curation and *a priori* estimates of strain numbers in order to achieve optimal results^12^. An intuitive and automated method to generate lineage-resolved complete MAGs is needed for analysis of more complex metagenome communities in order to reduce the time required to validate results.

Recent improvements in long-read sequencing technologies (such as Oxford Nanopore or Pacific Biosciences) have dramatically improved the quality of de novo genome assemblies of large eukaryotic genomes^16,17^. However, due to the high error rate of long error-prone reads, assembly algorithms still fail to disambiguate between highly similar sequences, such as segmental duplications in the human genome^18^. The recent development of highly accurate HiFi reads from circular consensus sequencing (CCS) on the Pacific Biosciences platform resulted in long accurate reads with error rates below 1% across the length of the read^19^ providing opportunity to improve assembly quality^20^ and even resolve both haplotypes of diploid genomes^21,22^. Recent attempts to sequence and assemble metagenomes with long error-prone reads^23–25^ or linked reads^3^ have resulted in few successes. While it has been demonstrated that long error-prone reads result in longer contigs than short reads^23,24^, the assembly of nearly all members of the community in singular circular contigs has still been elusive. Much of this may be due to the imperfect “length versus error-rate” trade-off^26^. Although the recently introduced metaFlye assembler improved reconstruction of complex environmental metagenomes using long reads^5^, it subsequently produced collapsed representations of similar bacterial strains.

Long and accurate HiFi reads have recently resulted in the first complete human genome assembly by the Telomere-to-Telomere Consortium and opened the era of “complete (T2T) genomics^17^”. Thus, this new technology could be suitable for traversing and assembling the highly repetitive orthologous genomic features present in metagenomes into lineage-resolved complete MAGs, enabling a new era of “complete metagenomics”. Furthermore, variant calling using HiFi reads has the added benefit of providing single molecule, physical evidence of sequence variant linkage that can be used to define discrete haplotypes in MAGs. In this paper, we leverage the use of HiFi reads in a metagenome assembly and demonstrate that metaFlye assembly with HiFi reads produce more lineage-resolved complete MAGs compared to assemblies generated with uncorrected long-reads without the need for manual curation. Additionally, we present a computational approach to phase alternative SNP haplotypes in these MAGs to provide finer resolution of descendant lineage variation in the sample.

## Results

### Assembly of the sheep gut microbiome

We extracted high molecular weight DNA from a fecal sample of an adult sheep collected during necropsy to determine cause of death. The resultant DNA prep was sequenced using a short-read (Illumina, San Diego, CA) and a long-read (PacBio, Menlo Park, CA) sequencing technology, with the latter using the CCS method to generate HiFi reads from the error-prone subreads (for a technical definition, see Methods). The short and HiFi reads comprised 154 and 255 total Gigabases (Gbp) in 1,024,375,790 and 22,118,393 reads, respectively, with the latter representing higher depth of coverage compared to most previous reports of long-read metagenome assembly. metaFlye assembly of HiFi reads resulted in a total of 57,259 contigs with a contig N50 of 279 kb, including 127 contigs that fit the criteria of a high-quality draft^1^ (or by our terms, “complete”) MAG. Among the MAG-quality contigs, 44 (35%) represented closed circles in the metagenome assembly graph (see Table 1).

**1.** Assembly quality statistics.

1. Assembly quality statistics
| Assembly | Contigs | Assembly Length (Mbp) | Contig N50 (Kbp) | HQ Draft Contigs<sup>1</sup> | Circular Contigs<sup>2</sup> | Circular + HQ Draft Contigs |
| --- | --- | --- | --- | --- | --- | --- |
| HiFi | 57,259 | 3,424 | 280 | 123 | 49 | 44 |
| pCLR1 | 48,338 | 2,985 | 185 | 54 | 21 | 18 |
| pCLR2 | 48,790 | 3,008 | 187 | 65 | 28 | 26 |
| pCLR3 | 56,456 | 2,978 | 181 | 64 | 26 | 22 |

<sup>1</sup> Contigs that were determined to have greater than 90% single copy gene (SCG) completeness and less than 5% SCG redundancy.
<sup>2</sup> Contigs larger than 1 Mbp in size that were predicted to be circular by the metaFlye assembler.

### Long accurate reads result in significantly improved metagenome assemblies

We hypothesized that substantial improvements in assembled contig completeness statistics were primarily due to the lower error rates of reads providing less ambiguity in resolving structural complexity in microbial genomes, so we sought to create an experimental design that would quantify the benefits of using an equivalent amount of long error-prone reads. Comparisons of metagenome assemblies based on generation of separate library types from the same sample may suffer from differences in microbial composition, the temporal nature of samples and the likelihood of sampling particular microbes in the community. As such, comparisons of a separate library of long error-prone continuous long reads (CLR) taken from the same DNA sample are unlikely to control for all of the confounding variables that impact the quality of the downstream assembly.

We devised an approach for an apples-to-apples comparison of HiFi and CLR reads by extracting subreads from the original HiFi reads to generate a series of “pseudo-CLR” (pCLR) datasets. We generated three separate assemblies corresponding to the first (pCLR1), second (pCLR2) and third (pCLR3) full-length sub-read, respectively (Figure 1a). These subreads share the same error profile as PacBio CLR sequencing (8-15% error rate^27^) but were equivalent in length to their parental HiFi reads. We assembled these datasets using metaFlye and compared them to our HiFi assembly to quantify the benefits of using long and accurate reads instead of long error-prone reads. The average pCLR contig was longer than the average HiFi contig in all Superkingdoms except the Eukaryotes (Figure 1b). However, the total assembly length of pCLR contigs was lower than the HiFi assembly in all categories except the unassigned, “no-hit” lineage (Figure 1c). In the Archaea and Bacteria annotated contigs, the pCLR assemblies had an average of 61 high-quality draft genomes with an average of 22 predicted circular complete genomes, representing a 48% and 50% reduction, respectively, compared to the HiFi assembly (Table 1; Figure 1d).

**Figure 1.**
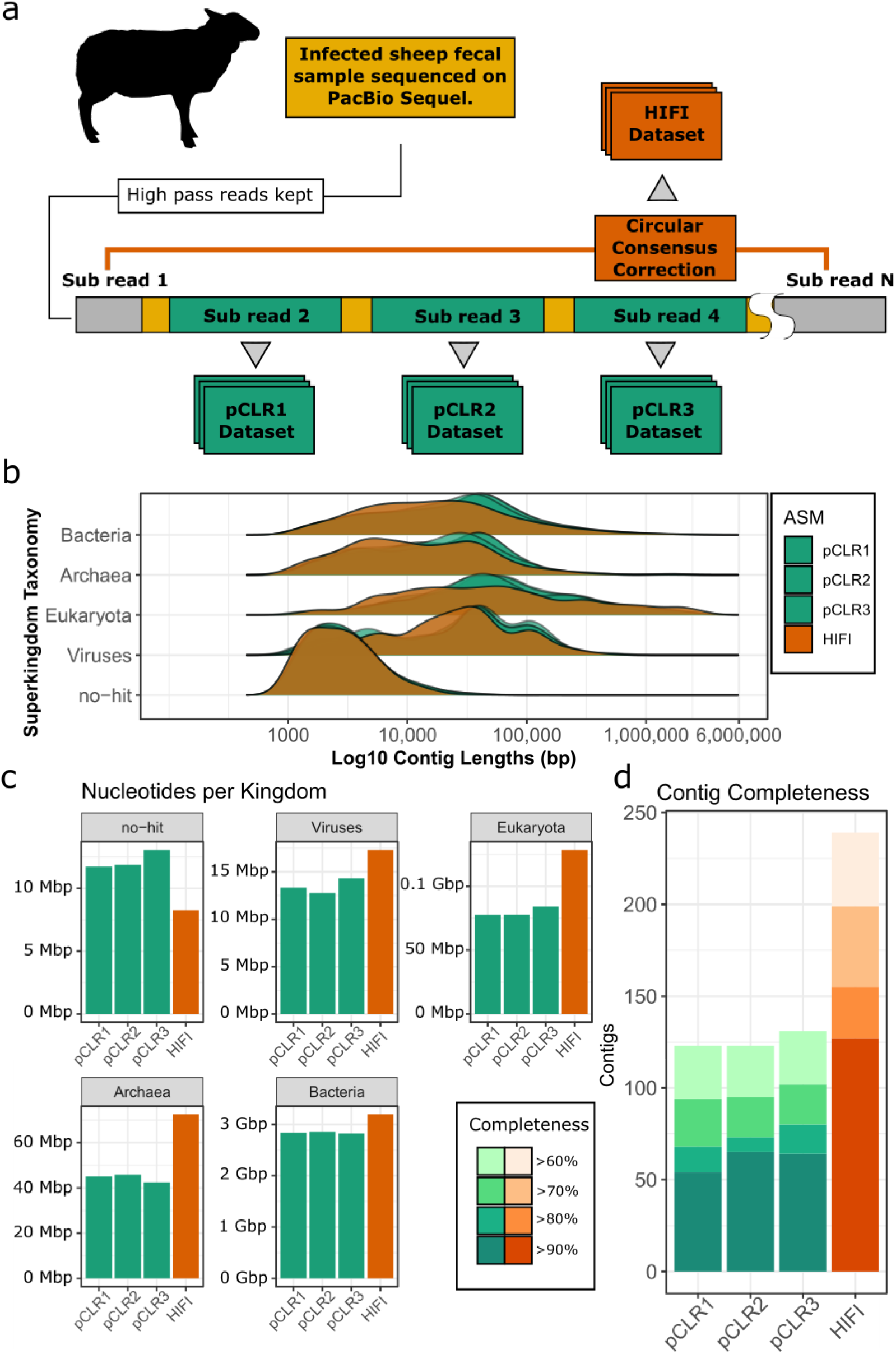
Contig-level comparison of pCLR and HiFi assemblies. a. Strategy for generating the read sets for the three pCLR and the HiFi assemblies. b. Comparison of contig length distributions in the four assemblies demonstrating a tendency for pCLR assembly to create longer contigs. c. Comparison of the total length of each assembly after separation of contigs into predicted Superkingdoms demonstrating an increased length from HiFi assembly among assigned Superkingdom and reduced length in unassigned bin. d. Comparison of the completeness of pCLR and HiFi assemblies based on the presence of >90% expected single-copy genes with <5% redundancy.

Binning the HiFi contigs with Hi-C linkage data (see Methods) resulted in 428 complete MAGs (> 90% SCG completeness and < 10% SCG contamination), which is the largest number of reference-quality MAGs reported from a single sample, to our knowledge (Supplementary Table 1). The pCLR assemblies also resulted in a substantial quantity of complete MAGs, with an average of 335 MAGs in each assembly (78% of the HiFi total). We hypothesized that one factor contributing to the lower number of MAGs in the pCLR assemblies could be a smaller number of contigs from distinct, related lineages that were more correctly represented in the HiFi assembly. Consistent with this hypothesis, a cumulative assembly length plot suggested that a larger proportion of complete MAGs in the HiFi dataset were of low relative abundance (with coverage below 10x) compared to MAGs in the pCLR assemblies (Figure 2a). Comparisons of bin SCG completeness and average depth of coverage also indicated that the HiFi assembly had more low-coverage complete MAGs than the pCLR assemblies (Figure 2b). The contrast between HiFi and pCLR assemblies was more pronounced in bins that had > 90% SCG completeness (Figure 2c), where the pCLR assemblies contained mainly bins with more than 10X coverage and as much as 1000X coverage compared to the HiFi complete MAGs. The distribution of coverage for complete MAGs is consistent with the hypothesis that HiFi assembly resolved pCLR bins into higher resolution, lower coverage bins that had been compressed into single bins in the pCLR assembly.

**Figure 2.**
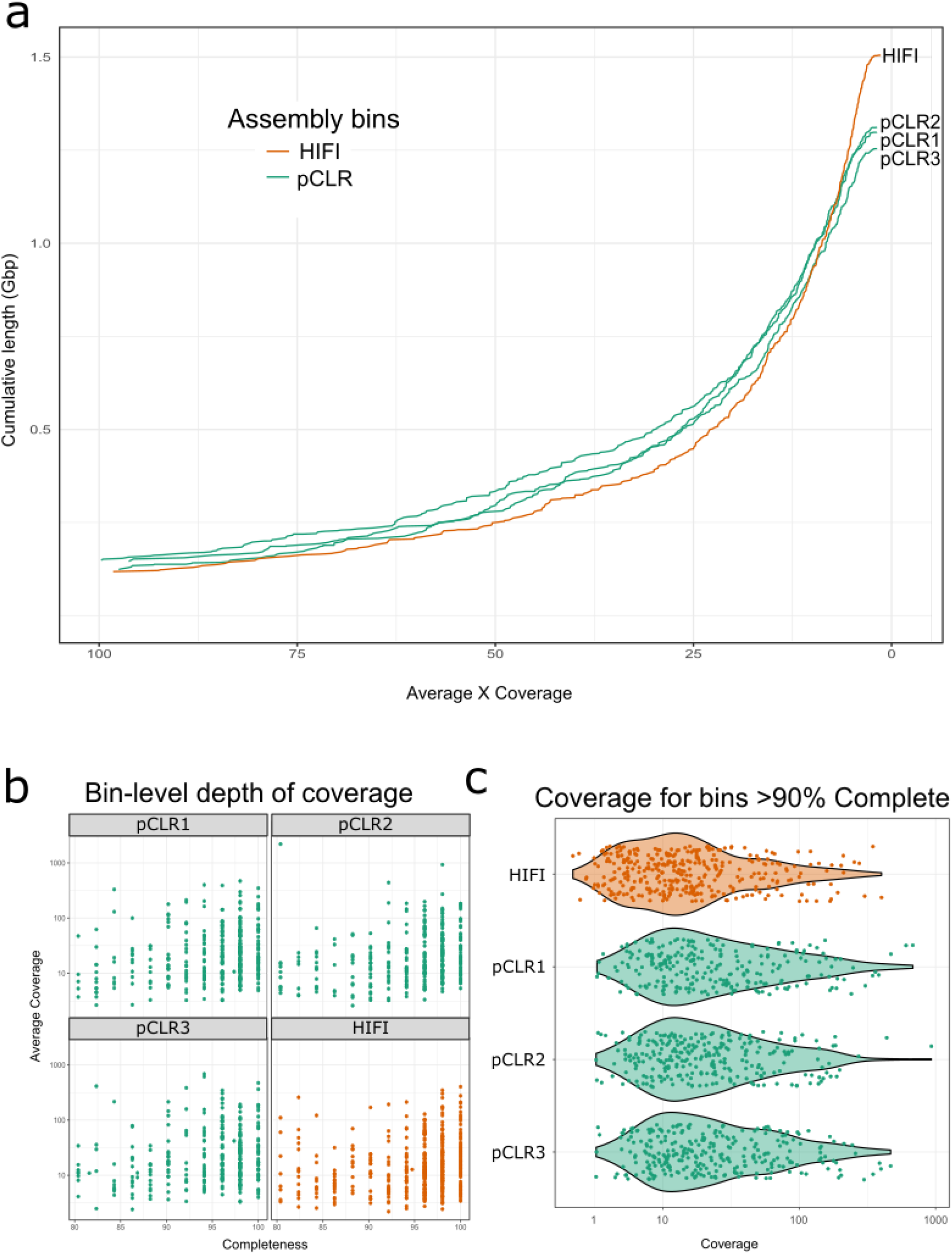
The cumulative length of assembled HiFi bins (a) peaks at lower depths of coverage at a faster rate than the cumulative lengths of pCLR bins, suggesting that lower abundance taxa were more likely to be assembled by HiFi reads. Comparisons of average short-read coverage against single copy gene completeness estimates (b) for high-quality bins revealed a substantial number of HiFi bins below the 10X coverage threshold compared to the pCLR datasets. This is particularly enhanced in the >90% completion category, where the average coverage of the HiFi bins is lower than that of each pCLR assembly, and several HiFi bins have less than 1X average short-read coverage as opposed to no equivalent coverage-profile pCLR bins.

### Lineage-resolved MAGs enabled by assembly with HiFi reads

Our experimental design allowed us to test the hypothesis that assembly with HiFi reads had separated distinct lineages into individual assemblies within metagenomes compared to assemblies with pCLR reads. We first classified HiFi and pCLR complete MAGs into predicted phylogeny using GTDB-TK^28^, resulting in 197 and 187 distinct Genera, and 15 and 14 distinct Phyla classifications, respectively (Supplementary Figure 3). There were 22 genera unique to the HiFi dataset, compared to 8 among all three pCLR datasets, and one phylum unique to HiFi bins (Supplementary Figures 4,5 and 6). Several cases where the HiFi assembly had more assembled bins for a taxon than the pCLR assemblies were also identified (Supplementary Table 2). A clear example of this was for a lineage assigned to the Clostridia class, which had three assembled bins in the HiFi assembly. These three bins had estimated MASH^29^ distance scores between 0.05-0.07, suggesting that they are separate assemblies of related organisms within this class and possibly represent different species within genus or strains within species (Supplementary Tables 3 and 4). Comparisons of alignments of contigs to the assembly graphs for these bins show clear separation of MAGs within the HiFi dataset and comparatively heterogeneous regions of alignment in equivalent, collapsed pCLR MAGs (Figure 3a). Separation of these HiFi complete MAGs was further compared to the three pCLR datasets through MASH kmer profile comparisons, which revealed that only one bin per pCLR assembly fell within a predicted MASH distance of 0.10 from any of the three HiFi MAGs. This suggested that the pCLR assemblies had collapsed the distinct components of the separate HiFi MAGs into single bins. Indeed, the pCLR contig bins corresponding to the Clostridia class had uneven depths of coverage averaging approximately 45-fold, suggesting they represent composites of distinct lineages, compared to consistent coverage across the contigs for the HiFi bins (Figure 3b). Moreover, this consistent read depth in the HiFi bins varied with the three bins having approximately 10x, 20x, and 33x coverage, demonstrating the potential to accurately deconstruct subtypes across a range of relative abundance. This outcome has significant implications in the use of read coverage in resolving strain lineages from metagenomes.

**Figure 3.**
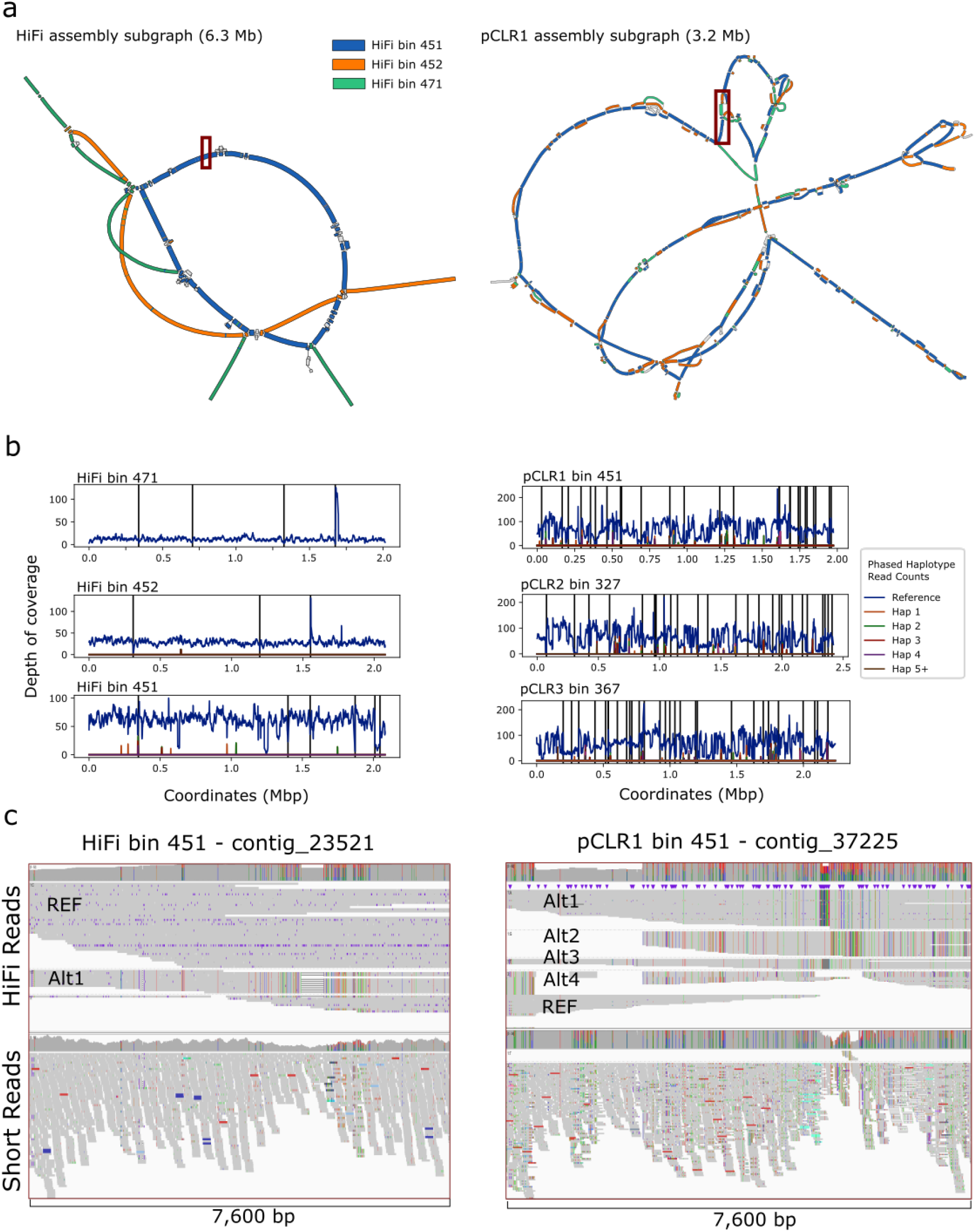
Lineage resolved MAGs (a) in the HiFi assembly often corresponded to two or more compressed bins in the pCLR assemblies. In this example, we show comparative alignments of HiFi bins (colored according to the legend) on superset graphs of three HiFi bins (left-most graph) and a single pCLR1 bin (right-most graph). pCLR graph alignments show bifurcation and trifurcation of sequence into bubbles that were otherwise condensed in the final assembly. Dark red tinted boxes correspond to IGV plots in (c). Using our newly developed MAGPhase algorithm (b), we identified several locations where multiple SNP-derived haplotype alleles are present in each bin (alternating colors) and estimated their relative depth compared to all HiFi read alignments. IGV plots of specific loci within these bins (c) show the power of this method to easily distinguish between haplotypes without the need for extensive statistical post-hoc analysis. Comparative alignments of HiFi reads to HiFi bin 451 show only one alternate allele, whereas the equivalent region in the pCLR1 bin 451 shows as many as four alternate alleles (labeled on the figures). Furthermore, comparisons with short-read alignments revealed the inadequacy of short-reads to identify phased haplotypes within these highly resolved MAGs.

The Clostridia class was instructional but was not the only example of collapsed assemblies present in the pCLR MAGs. A total of 15, 10 and 11 pCLR MAGs were found to be condensed orthologs of 31, 23 and 25 HiFi bins in the pCLR1-3 assemblies, respectively (Supplementary Figures 7, 8 and 9). We also identified other MAGs within the HiFi assembly that are likely species- or strain-resolved assemblies using a nearest neighbor distance analysis with a low MASH pairwise distance cutoff (<= 0.07 distance). These MAGs likely represent “lineage-resolved” assemblies of individual subpopulations within the same sample as they are separate assemblies of organisms from the same genus or species given this distance cutoff. We identified 18 such MAGs within the HiFi assembly, which was triple the amount in the pCLR assemblies (an average of six lineage-resolved MAGs; Supplementary Table 5). These HiFi MAGs had solitary representatives in the pCLR assemblies, suggesting that such fine-scale differences in sequence content and structural variation are likely to be lost in assemblies of long error-prone reads.

### Improving resolution within lineage-resolved complete MAGs using HiFi reads

Comparison of pCLR and HiFi bins demonstrated that HiFi assembly resolves sub-lineages even at the stage of initial contig output from the metaFlye assembler. This result motivated us to investigate whether we could further resolve HiFi bins into lineage-resolved complete MAGs using SNP variant data as attempted previously^14^. We identified several MAGs that still had single nucleotide polymorphism (SNP) variation above that expected from read error rates within SCG regions. Alignments of short-reads were unable to distinguish true polymorphic sites, particularly in highly repetitive or frequently orthologous gene regions (Figure 3c), so we developed a computational approach to resolve lineages in metagenomes. This approach required an ability to distinguish between polymorphisms within a lineage and structurally variant subtypes within a MAG, which in turn required an ability to simultaneously consider depth of coverage and haplotype information.

Since this problem has similarities to phasing isoforms of transcripts in the context of variable expression from parental alleles in gene expression studies, we adapted the phasing algorithm of the IsoPhase workflow^30,31^ into a new tool called MAGPhase to identify SNPs on individual HiFi reads and to phase them across identified single copy gene regions in each MAG. To avoid potential false positive SNP haplotypes due to errors in reads, we only call variants in SCG regions that have at least 10 spanning HiFi reads, and are prevalent at significant proportions of read depth as assessed by a Fisher exact test with Benjamini-Hochberg^32^ correction (see Methods). Phased SNP haplotypes were identified in each target region and the maximum number of haplotype alleles was counted for each MAG to assess the upper boundary for SCG variation in each MAG. A majority of HiFi MAGs (220; 52% of the total) had zero identified alternate haplotype alleles, suggesting that many lineages were well resolved by the HiFi assembly or did not have detectable polymorphic subpopulations in the sample (Table 2). This is in contrast to the pCLR assemblies, of which an average of 118 MAGs (35% of the total) were found to have zero haplotype alleles (Supplementary Table 6). Polymorphic HiFi MAGs were found to exhibit as many as 25 unique haplotype alleles within SCG regions, suggesting localized regions of genetic drift. This is further supported by the fact that, among 48 HiFi haplotype loci with more than 10 unique alleles, we found that 40% (122/305 haplotypes) differed from the original reference sequence by three or fewer bases, suggesting fixation of neutral mutations in subpopulations^33^. Median coverage of the alternative alleles in these hotspot regions was an average of five HiFi reads across the length of the haplotype, suggesting that most of these alternative haplotypes were likely not caused by read errors.

**2.** MagPhase Haplotyping results.

2. MagPhase Haplotyping results
|  | HiFi | pCLR1 | pCLR2 | pCLR3 |
| --- | --- | --- | --- | --- |
| Total complete MAGs | 428 | 345 | 345 | 315 |
| Average Contig count per MAGs | 8.3 | 10.8 | 11.4 | 11.9 |
| Zero Haplotype MAGs<sup>1</sup> | 220 | 130 | 136 | 89 |
| Percent Polymorphic MAGs<sup>2</sup> | 48.5% | 62.3% | 60.6% | 71.7% |
| Average Haplotype Variant Length (bp)<sup>3</sup> | 20.1 | 27.1 | 24.1 | 34.0 |
| Average Haplotype Genomic Length (bp)<sup>4</sup> | 1151.3 | 1106.1 | 1057.8 | 961.2 |
| Maximum Haplotype Genomic Length (bp) | 336,899 | 463,082 | 480,257 | 493,333 |
| Average Haplotype Alleles per Locus<sup>5</sup> | 4.18 | 4.72 | 4.43 | 5.06 |
| Maximum Haplotype Alleles per Locus | 25 | 59 | 54 | 60 |

<sup>1</sup> Metagenome assembled genomes (MAG) that did not have detectable SNP haplotypes that could be linked with HiFi reads.
<sup>2</sup> The number of MAGs that had at least one detectable alternative SNP haplotype allele.
<sup>3</sup> Haplotype variant length represents the count of polymorphic SNP loci within an identified haplotype.
<sup>4</sup> Genomic length was defined as the distance in bases on the assembled contig from the first polymorphic SNP site to the final site.
<sup>5</sup> A haplotype allele was defined as a unique permutation of polymorphic SNP variants that were identified in one consistent region in the genome that met thresholds for detection.

Comparisons of aligned short-reads to polymorphic HiFi MAGs revealed limitations in the use of short-reads for strain heterogeneity assessments. Using the previously identified example of the lineage-resolved Clostridia MAGs, we identified 7, 1 and 0 alternative haplotype loci on HiFi bins 451, 452, and 471 respectively (Figure 3b). Closer examination of these regions revealed clear variant patterns in individual, aligned HiFi reads, demonstrating the power of using these data for phasing haplotypes from metagenome bins (Figure 3b). These signatures were not readily apparent or were heavily fragmented in the short-read alignments to the HiFi bins (Figure 3c). Furthermore, read pileups in lineage-resolved complete HiFi MAGs and orthologous pCLR collapsed MAGs were instructional in determining how read mapping could be used in downstream variant calling workflows. Comparing orthologous regions between the HiFi MAG 451 and pCLR1 MAG 451 (the similarity in number was a coincidence), the visual determination of haplotypes within the selected HiFi MAG is trivial (Figure 3c). One haplotype lineage containing a large insertion of sequence is clearly visible from read pileups and is identified by MAGPhase. By contrast, the pCLR1 MAG has four distinguishable haplotype alleles, consistent with the properties of a collapsed assembly. HiFi MAG 451 can consequently be separated into two separate lineage-resolved complete MAGs using these identified haplotypes, whereas the pCLR MAG is more difficult to resolve. In addition to this example, we identified 35 and 32 complete HiFi MAGs that had only 1 or 2 identified alternative SNP haplotypes that could be separated into an additional 70 and 96 lineage-resolved complete MAGs, respectively. However, we note that 220 of our complete MAGs had zero identified haplotypes without any need for manual curation, and therefore fit the criteria of lineage-resolved complete MAGs by default. We adopt this tally of 220 lineage-resolved complete MAGs as a final, conservative estimate of metaFlye assembly of our HiFi read dataset to demonstrate the lack of need for extensive post-hoc editing. In both assemblies, short-read alignments also failed to consistently identify variants within identified haplotype alleles, regardless of the quality of the underlying MAG.

The paucity of consistent signal and the smaller power to link variants into haplotypes appears to limit the use of short-reads for variant phasing in complex metagenome communities. Furthermore, the prevalence of many ambiguous short-read alignments with a mapping quality score of 0 (MapQ0) in haplotype regions suggests that these regions are highly repetitive in the overall assembly and do not provide sufficient unique sequence for short-read alignment. The percentage of short-read MapQ0 alignments out of the total were 7%, 9% and 17% for bins 451, 452 and 471, respectively, suggesting that large portions of these bins would be otherwise intractable to variant profiling using short-read data. Indeed, a 5 kb window analysis across the entirety of the HiFi assembly identified 18% of the assembly is covered by windows that have ratios of MapQ0 alignments to total alignments greater than 0.50. Naturally occurring variation is unlikely to be detected in these windows via short-read alignments due to mapping ambiguity. By contrast, the proportion of high MapQ0 HiFi alignment windows were found to constitute only ∼ 2% of the length of the assembly, suggesting that 98% of the assembly contains sufficient unique sequence for HiFi read alignment (Supplementary Figure 10).

### Improvements in functional genetics analysis

We illustrate the advantages of HiFi reads in functional annotation of a metagenome by predicting biosynthetic gene clusters (BGCs) that are notoriously difficult to identify in fragmented assemblies^34^. We identified 1,400 complete and 350 partial BGCs (the latter being defined as lying on a contig edge) in the HiFi assembly using antiSMASH^35^. To the best of our knowledge, this represented the largest number of complete BCGs ever reported in metagenomic assemblies and a 40% increase over discovery rates in the three pCLR assemblies (showing 1,245 BGCs on average), with appreciable increases in the detection of important nonribosomal peptide synthetase (NRPS, 25 %) and ribosomally synthesized and post-translationally modified peptide (RiPP, 40%) BGC classes (Fig. 4a). This substantial increase in detected BGCs is not commensurate with the increase in assembly size (15% more assembled HiFi sequence), suggesting that the BGC prediction was significantly improved in the HiFi assembly. Interestingly, nearly all identified BGCs were classified as novel, in the sense that no reference gene clusters of known function were found with >50% of their genes showing homology, illustrating the capabilities of long reads for exploration of novel natural products. We identified 40% more novel BGCs in the HiFi assembly than in the pCLR assemblies (Fig. 4b). Finally, more partial BGCs were identified in the HiFi assembly (Fig. 4c).

**Figure 4.**
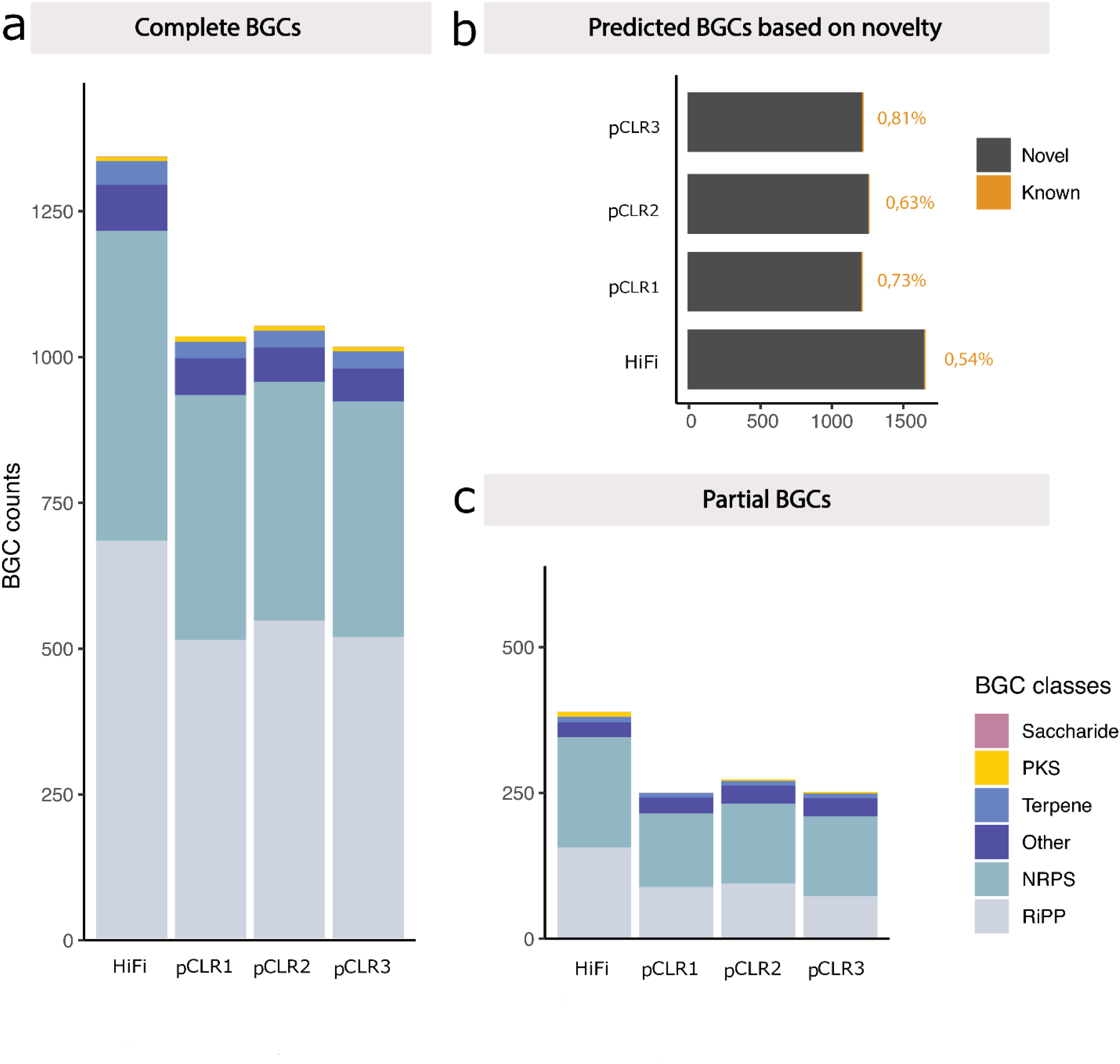
The HiFi assembly revealed approximately 25% more complete Biosynthetic Gene Clusters (BGCs) than the average pCLR assembly (a). This increase was manifested in all identified BGC classes (colors in legend) and was not exclusive to one particular class. As found in other metagenome assembly datasets, the majority of identified BGCs were novel in all assemblies (b), but the HiFi assembly had a higher proportion of novel BGCs than the other assemblies. Additionally, the HiFi assembly contained more partial BGCs (c) of any assembly.

### Improved resolution of mobile DNA association analysis

Candidate viral contigs were identified in each assembly using an alignment-based approach (see Methods). Candidate viral contigs ranging from 5-250 kb in length were identified in each assembly using an alignment-based approach (see Methods). The higher count of assembled viral contigs in the HiFi assembly (*n* = 383) compared to the pCLR assemblies (average: 276; stdev: 20) suggested that the breadth of viral diversity in the sample was best represented in that assembly. We conducted an association analysis of viral contigs to candidate microbial hosts using Hi-C links and partial long-read alignments by application of a previously published workflow^23^. A resulting network analysis showed that the majority of the viral associations were found between the viral *Siphoroviridae* family and bacterial hosts (Fig 5a), regardless of the use of the HiFi assembly (211 associations), or the pCLR assemblies (185.7 average associations; Supplementary Figures 11, 12 and 13). More associations due to long-read overlaps were identified in the HiFi viral network than the pCLR network (Fig 5b), likely due to improved alignment mapping rates in that assembly. Interestingly, the HiFi assembly provided more evidence of Virus-Archaea associations (60 archaeal contigs) than the pCLR datasets (8.5 mean contigs), primarily via partial long-read alignment metrics which are evidence of genomic integration via a lysogenic life-cycle phase^23^(Fig 5c). This increase in Archaea-viral associations is likely due to the increased assembly of Archaea-origin sequence in the HiFi assembly (Figure 1c), which enabled the detection of integrated archaeal virus sequence.

**Figure 5.**
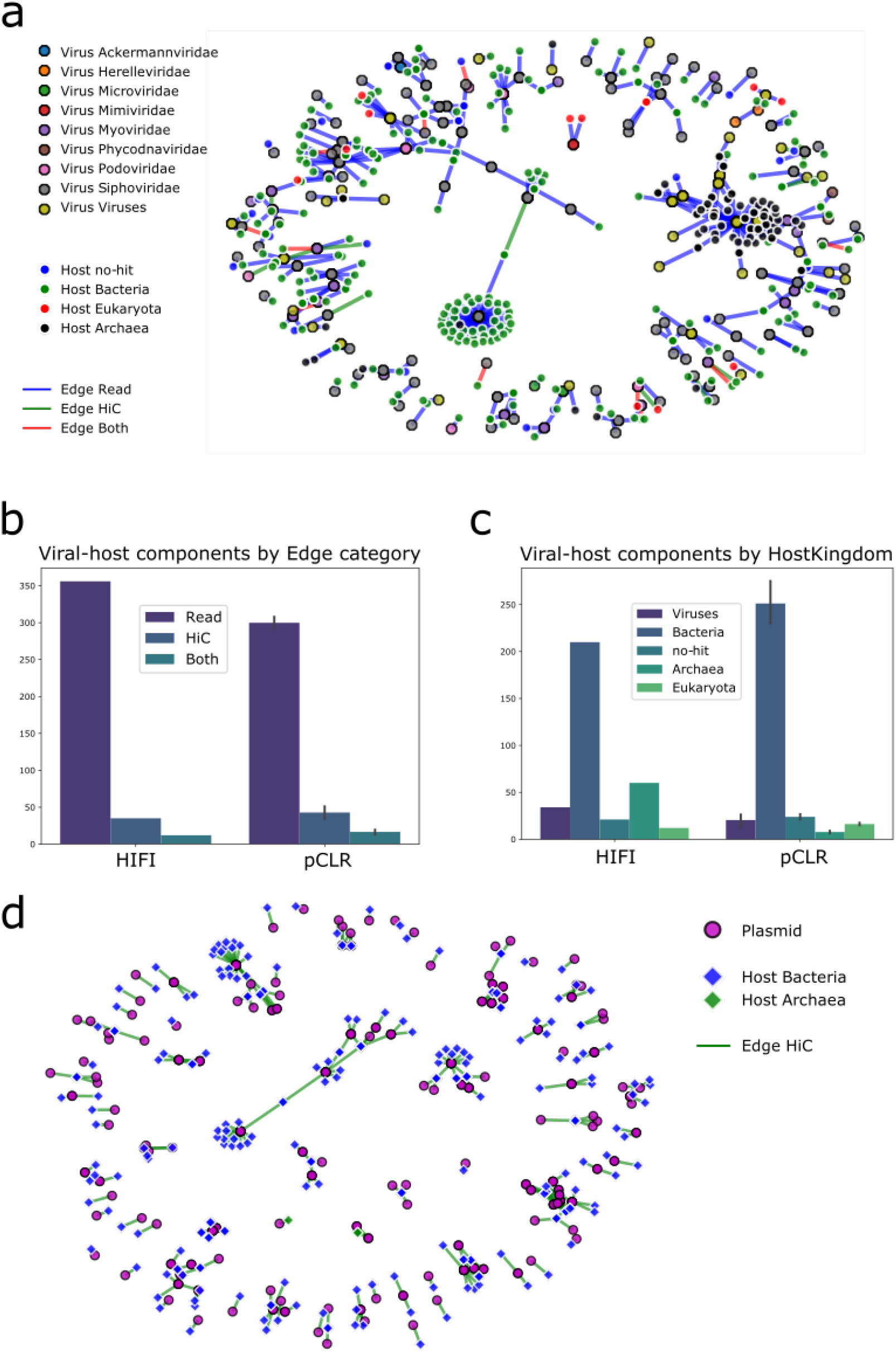
A network plot of predicted host-virus associations (a) identified through HiFi read overlaps (blue), Hi-C links (green) and both data types (red) revealed new viral genomes that have broad host specificity. In addition, the HiFi assembly was better able to identify candidate viral-Archaeal associations than those detected in the pCLR datasets. Viral-host associations were predominantly identified through HiFi read alignments (b) and the HiFi assembly had a higher proportion of this evidence compared to the average pCLR assembly. Highlighting the difference in domain detection between the assemblies, more Viral-Archaeal links (c) were identified in the HiFi assembly compared to the pCLR assemblies. Using Hi-C link data, we were also able to identify candidate hosts for assembled plasmid sequence (d) in the HiFi assembly.

Our HiFi assembly also contained many predicted short circular contigs (< 1 Mbp) that likely represented complete plasmid sequence. Using the SCAPP^36^ plasmid assembly tool, we identified 5,528 candidate plasmid contigs within the HiFi assembly. We identified 298 plasmid-contig associations in the HiFi dataset using Hi-C linkage data (Fig 5d). The largest subgraph (degree = 18) consisted of an interesting association between six plasmid contigs and 25 candidate bacterial hosts (Supplementary Figure 14), in which one plasmid was predicted to inhabit members of 13 different bacterial genera, suggesting inter-genera mobility of this plasmid. We also predicted associations between identified plasmid contigs and three genera of Archaea, including *Methanobrevibacter* and *Methanosphaera*, which were previously not known to carry naturally-occurring plasmids^30^.

## Discussion

The goal of metagenome assembly is to create representative reference genomes for the majority of organisms that comprise the sample. However, our data suggests that both short and long error-prone reads produce collapsed assemblies that would otherwise require extensive manual curation to resolve into reference-quality resources. Here, we show that metaFlye assemblies using HiFi reads generate lineage-resolved complete MAGs for single samples without the need for curation (Figure 3). Furthermore, we found good representation of organisms that tend to be prevalent at lower relative abundance in the community in assembled MAGs (Figure 2a), but we nevertheless assembled them to meet the criteria for high-quality draft genomes^1^ (Figure 2c). These complete MAGs appear to be resolved with respect to structural variation and orthologous gene sequence compared to closely related (< 10% MASH distance) lineages as evidenced by the assembly graph comparisons. Our data suggests that lineage-resolved complete MAGs are difficult to generate using long error-prone reads, and our experimental design shows that the accuracy of HiFi reads is a necessary element of this result. Sketch-based comparisons revealed that several of the HiFi MAGs (23 - 31 MAGs; 6 - 7% of total) were condensed into collapsed assemblies in the pCLR datasets. The collapsed pCLR bins present in the pCLR dataset were found to be poor representatives of the actual genomic sequence of the organisms based on read alignment metrics (Figure 3b) and variant phasing analysis (Figure 3c). This may present a future challenge for other long-read metagenome assemblies, as lineage-resolved MAGs are most likely to be collapsed in such surveys, particularly if several closely-related species are present in the sample.

Our variant phasing method with HiFi reads greatly simplifies variant lineage detection within a sample through the detection of discrete haplotypes. Existing short-read-based strain-resolution algorithms rely on multiple sample observations and statistical variant linkage analysis in order to determine potential microbial lineages^11,37^. By contrast, HiFi reads provide suitable accuracy and length to enable easy identification of linked variants within a single sample. We identified phased haplotypes of up to 309 SNPs and phase variants across segments as large as 300 kbp in our HiFi MAGs (Table 2). Rather than limiting analysis of microbial lineages to average nucleotide ID (ANI) thresholds that may be biased due to short-read alignment inaccuracy, HiFi reads allow for detection of haplotypes segregating in a sample that have as low as 2% (5 reads out of 300) relative abundance of the reference MAG haplotype. To enable this degree of classification, we provide a pipeline and workflow called MAGPhase, based on the cDNA_Cupcake API that is the first to use HiFi reads for haplotype analysis on metagenome assemblies (https://github.com/Magdoll/cDNA_Cupcake). Our IGV alignment diagrams show that evidence supporting the prevalence of these SNP haplotypes is visually verifiable due to the accuracy of HiFi reads. We provide tools to reproduce these diagrams within the MAGPhase workflow to assist future surveys using this data. Even when using MAGs produced by long error-prone reads (pCLR assemblies) as a reference, MAGPhase can still produce discernable SNP haplotypes that could be used to identify descendant lineages (Figure 3c). This means that existing references from isolates or other long-read metagenome surveys could be used in tandem with HiFi reads for strain-typing. However, we note that HiFi alignments to lower quality MAGs are likely to contain far more noise than when using lineage-resolved complete MAGs as references, so *de novo* HiFi-based assemblies are still preferred in this context.

We acquired several biological insights from our data that were provided almost exclusively by the HiFi assembly and HiFi reads. Use of the antiSMASH^35^ detection tool identified 40% more BGCs in the HiFi assembly than the highest count in the next best pCLR assembly. The antiSMASH results also provided insights into the functional potential of secondary metabolic pathways in the sheep gastrointestinal tract; for example, 19 BGCs were found in the HiFi data that show high similarity to a recently identified class of gene clusters encoding the production of proteasome inhibitors from the human gut microbiota^38^, indicating that these functions may be of similar importance for host colonisation in ruminants as they are in humans. More of these BGCs were predicted to be novel in the HiFi assembly, and were furthermore not found to be resolved replicates of compressed consensus sequences in the pCLR assemblies. Additionally, we identified several novel associations of mobile genetic elements in our sample using a combination of Hi-C linkage data and HiFi read alignment overlaps. Both the pCLR assemblies and HiFi assembly had similar profiles of viral-host association links, with a notable exception in the case of links between Archaea and viral contigs. The HiFi assembly detected a higher quantity (n= 60) and greater complexity (diameter = 7) of archaeal-viral associations primarily through HiFi read overlaps. Host-plasmid analysis using Hi-C links also identified broad host-specificity for several assembled, circular plasmids in our HiFi dataset. In total, we identified 424 and 298 potential host-viral and host-plasmid links in our HiFi dataset, which represents one of the most substantial associations of mobile element activity in a single sample to date. Most of these associations were exclusive to the HiFi assembly and were not identified in replicate pCLR assemblies.

The improvements in lineage resolution and haplotype phasing offered by HiFi reads present new opportunities but also a major dilemma. HiFi reads are currently more expensive to generate than equivalent amounts of short-reads and long error-prone reads. Additionally, HiFi reads tend to be shorter than reads generated by PacBio CLR mode or Oxford Nanopore platforms due to shorter molecule fragment size requirements for circular consensus sequencing (CCS) to obtain enough passes across the insert (minimum of 3) for CCS error correction. This could limit their application in large-scale metagenomic surveys; however, we note that DNA fragment size distributions from recent long-read metagenome assembly surveys often do not exceed 10 kbp in size^23,24^. In the absence of a reliable protocol to generate metagenome WGS datasets with read N50 values above 100 kbp as per typical “ultra-long” library preparations^17^, the choice between longer CLR datasets and higher quality HiFi datasets could be a false dilemma. We previously reported that the use of long error-prone reads resulted in a four-fold increase in contig N100K statistics over a comparable short-read assembly^23^. In this study, we found that the use of the same amount of near-equivalent length, HiFi long-reads resulted in a 2.5-fold increase in SCG complete contigs over assemblies constructed from their constitutive subreads. Another HiFi-specific advantage is the assembly of lineage-resolved MAGs that were otherwise condensed in pCLR assemblies, and the phasing of variant haplotypes to distinguish finer resolution differences in populations in the sample.

To our knowledge, this is the first time that it has been possible to examine the population structure of metagenomes using whole assemblies and read-phased haplotype alleles, and it creates more exciting possibilities for future study. Our analysis suggests that such insights can only be gained through the use of long (> 5 kb) reads with suitably low (∼ 1%) error rates, as the former criteria enables the spanning of orthologous genomic regions and the latter enables the separation of species/strain-level haplotypes into separate assemblies. These results were obtained with relatively minimal efforts, requiring only assembly and binning of HiFi reads with Hi-C data, thereby obviating the need for extensive manual curation. Resulting lineage-resolved complete MAGs and phased SNP haplotypes are the first realization of “complete metagenomics” - isolate-quality genome assemblies for microbial organisms from complex metagenome samples.

## Online Methods

### Long-read DNA sequencing and subread extraction

A fecal sample was taken from a young (<1 year old) wether lamb of the Katahdin breed. The animal died while on pasture and postmortem was diagnosed with combined *Strongyloides* and coccidial infection. The sample was acquired postmortem following the USDA ARS IACUC protocol #137.0 during routine necropsy to determine cause of death. The sample had a watery texture consistent with diarrhea and apparent parasite eggs were observed within the sample, which was transferred to a 50ml tube, mixed to make as homogenous as possible, and aliquoted into 1.5 ml microfuge tubes. DNA was extracted in small batches from approximately 0.5g/batch using the QIAamp PowerFecal DNA kit as suggested by the manufacturer (QIAGEN) with moderate bead beating and sheared using a Digilab Genomic Solutions Hydroshear instrument (Digilab). The sheared DNA was size-selected to approximately 9-18 kb on a SAGE ELF instrument to final target size which varied from 9 kbp up to 16 kbp followed by library preparations using the SMRTbell Template Prep kit v1.0 as described (20). Sequence data was collected over time and included 46 SMRT cells on a Sequel instrument using 10 library preparations, with 24 cells of v2 chemistry and average inserts of 9-10 kbp and 22 cells of v3 chemistry and average inserts of 14 kbp. An additional 8 cells representing individual library preparations were sequenced on a Sequel II instrument using v1.0 chemistry and average inserts of 14 kbp. Subreads and CCS reads were generated using SMRTLink software v6.0 CCS protocol and default settings. An average of 35% of subreads per cell were converted to CCS corrected reads (range 1-63%). This resulted in 255 Gbp of total CCS reads from both the Sequel I (45 Gbp of the total) and Sequel II (210 Gbp) sequencing runs. A subset of this data (46 Sequel I SMRTcells) representing 18% (45 Gbp) of the total dataset was previously assembled as validation data in the metaFlye assembler publication^5^. The Sequel II dataset was filtered after CCS correction to retain only reads that fit HiFi quality standards (3+ full length passes and average read quality scores > Q20). We note that a small proportion of our Sequel I dataset (4,350 reads; 0.02% of the total number of CCS reads) consisted of reads that did not meet HiFi read quality standards (average Q scores above 20) as this dataset had been filtered with a prior version of the SMRTLink software. These reads were retained as they comprised a very small proportion of the total dataset.

Subreads were extracted from the converted CCS reads to provide a suitable comparison between uncorrected and corrected long read datasets. First, all of the constitutive subreads of the CCS reads were identified from subread BAM files. Using a custom script (https://github.com/njdbickhart/python_toolchain/blob/master/assembly/extractPacbioCLRFromCCSData.py), the second, third and fourth subreads were separately extracted into FASTQ files designated pCLR1, pCLR2, and pCLR3, respectively (the first subread does not typically encompass the complete DNA fragment, so was discarded). Statistics on subread lengths from the Sequel I and Sequel II datasets are shown in Supplementary Figures 1 and 2, respectively. Due to sequence read falloff in later subreads, the pCLR3 dataset was truncated relative to the pCLR1 and pCLR2 datasets. In the Sequel I dataset, a small proportion of reads (51 reads; ∼0.001%) did not have a fourth (pCLR3) subread, making the third replicate dataset smaller than the others. This resulted in a reduction of 104 Mbp of sequence in this dataset (48.763 Gbp) compared to the pCLR1 or pCLR2 extracted subreads (48.830 Gbp). The original CCS reads were organized into a dataset hereafter referred to as the “HiFi” reads and the three subread replicates were labelled pCLR1-3 in the chronological order in which they were sequenced in the subread BAM files.

### Short-read sequencing and Hi-C library preparation

An approximately 2g subsample of frozen homogenized fecal material was provided to Phase Genomics (Seattle, WA) for Hi-C contact map construction using their Proximeta service. The restriction endonucleases Sau3AI and MluCI were used to generate separate Hi-C sequencing libraries as previously described^39^. Using a total of 107 million paired-end reads from both Hi-C libraries were generated for analysis. A separate portion of the extracted DNA from the fecal sample was saved for short-read “whole genome shotgun” (WGS) DNA sequencing. Truseq PCR-free libraries were created from this sample as previously described^40^ and were sequenced on an Illumina NextSeq 500. A total of 149 Gbp of WGS short reads were generated from this sample.

### Genome assembly, read alignment and binning

Reads from the HiFi and pCLR datasets were assembled into contigs using the metaFlye^5^ genome assembler, version 2.7-b1646 for HiFi reads and version 2.7.1-b1590 for pCLR reads. The assembler was run in metagenome mode (“—meta”) flag and the “—pacbio-hifi” and “—pacbio-raw” data prefix flags were used for input HiFi reads and the pCLR reads, respectively. We note that the “—pacbio-hifi” input designation only uses reads that have average error rates below 1% for the disjointig and contig phases of the workflow. This means that only HiFi quality reads (Q20+) were used to generate the initial graphs and final contigs of the HiFi assembly. However, all input reads were used in the consensus polishing step of metaFlye. All assemblies were polished with two rounds of Pilon^41^ correction using the previously generated, WGS, short-read datasets. Contigs shorter than 1000 bp in all assemblies were removed from further analysis. Closed circular contigs were identified from metaFlye assembly reports. WGS short reads were aligned to the assemblies using BWA MEM^42^ using default settings. HiFi reads were aligned using Minimap2^43^ with the “-x asm 20” preset setting as recommended by the developers. Window-based alignment analysis was conducted by using custom python scripts (https://github.com/njdbickhart/python_toolchain/blob/master/sequenceData/getBAMMapQ0Ratios.py).

Hi-C read-pairs were aligned to each assembly using BWA MEM with the “-5SP” flag to disable attempts to pair reads according to normal Illumina paired-read settings. Resulting BAM files from Hi-C reads were sorted by read name. Hi-C alignments were used in the Bin3c^44^ binning pipeline to generate a set of bins for each assembly. Bin quality was assessed by CheckM^2^ and DASTool^45^ single copy gene metrics. MAGs were identified from bins that had > 90% SCG completeness and less than 10% SCG contamination estimates from the DASTool quality assessment data.

### Taxonomic assignment

We distinguish between contig-level and bin-level taxonomic classification to demonstrate differences in pCLR/HiFi assembly quality and assign representative taxonomy of the final polished bins, respectively. Contigs were assigned to candidate taxa using the Blobtools v1.0^45^ taxify pipeline, using models from the Uniprot (release: 2017_07) database as described previously^23^. Contigs that did not meet the Blobtools threshold for taxonomic assignment, or were identified as belonging to faulty database entries (e.g. the “Cetacean” lineage) were labeled as “no-hit” taxa. Viral contigs identified from this analysis were used in subsequent virus association analysis (see methods section below). Predicted viral contigs were separately verified using the CheckV^46^ pipeline using the “end-to-end” workflow, multithreaded, and with normal settings.

The GTDB-TK v1.0^28^ ‘classify_wf’ workflow was used to assign candidate taxonomic affiliation to all assembled bins. Default GTDB-TK settings were used with the only exception being the setting of the ‘—pplacer_cpus’ argument to ‘1’ as recommended by the authors. In cases where GTDB-TK was unable to assign a taxonomic lineage, a consensus of contig-level assignments from the Blobtools taxify pipeline were used to assign candidate taxonomic affiliation for the bin. The prevalence of three or more contigs in the MAG indicating the same species-level taxonomy were used when possible. In the case of “ties” between contig-level taxonomic consensus, the final taxonomic consensus was resolved to the lowest possible level (ie. genus or family).

### MagPhase lineage-resolution and orthologous MAG identification

We first sought to identify orthologous bins among each of the pCLR assemblies and the HiFi assembly in order to provide direct comparisons among similar assembled taxonomic groups. To identify orthologous bins, we used MASH v2.2^29^ sketches of all HiFi bins as a reference against queries of all pCLR bins. MASH sketch settings were -s 100000 and -k 21, with all other settings left at the default. The MASH “dist” command was used with a cutoff of 0.10 distance to identify orthologous MAGs which is approximately equivalent to an average nucleotide identity of 90% between hits. Multiple reference and query hits were allowed and retained for future comparisons.

HiFi reads were realigned to the HiFi and pCLR assembly bins using minimap2^43^ as previously described and alignment files were converted to BAM file format using Samtools^47^. To reduce the possibility of supplementary or split-read alignments impacting downstream variant calling, we filtered these alignments from the HiFi read BAM files. We then used these filtered alignment files for variant calling and haplotype identification using MAGPhase. The MAGPhase algorithm attempts to identify full-length SNP variant haplotypes in a greedy fashion within a given set of genomic coordinates. By default, only genomic coordinates that have at least 10 full length reads (10X coverage) are considered for variant calling. Initial variants used in read phasing are identified from HiFi read alignment pileups. To distinguish between potential errors in reads and SNPs, we model expected errors at a rate of 0.5% and test observed variant coverages against an expected error variant coverage using the Fisher exact test. To correct for further multiple hypothesis testing across an entire region, we employ a Benjamini-Hochberg^32^ procedure to estimate modified p values. The default p value cutoff for a variant site to be included is less than 0.10 for the modified p value. Once a set of candidate variants is called, the program then attempts to phase them into haplotypes based on their observed presence in HiFi reads. The entire length of an alternate haplotype is imputed using the physical linkage of previously identified SNP variants on individual HiFi reads that span the region. Missing variant information from imputation (due primarily from chimeric read alignments or the presence of read errors) is denoted with question marks (?) in the final haplotype dataset.

In order to reduce the potential expansion of haplotype counts due to recombination, we phased HiFi reads within identified SCG regions of each HiFi and pCLR bin. CCS reads that extended over the edges of SCG regions were included in haplotype phasing, so if two SCG regions were within short distances from each other, phased variant haplotypes could extend further. Partially imputed haplotypes (haplotypes that contained question marks (“?”)) were excluded from analysis as these could have resulted from chimeric read alignments or base call errors on selected SNP variant sites within the haplotype. Haplotypes were considered alternative alleles based on read depth, with lower depth haplotypes considered to be alternatives to the highest read depth allele at that loci. Haplotypes that included fewer than 3 SNPs were filtered as these tended to have lower counts of read alignments and higher alternate allele haplotype counts. If a MAG was found to have no SNP variants that fit the read depth statistical requirements, it was considered to be a “lineage-resolved” MAG. MAGs that had unfiltered SNP variants that were otherwise unable to be assigned to haplotypes with 3 or more SNPs were not considered to be lineage-resolved and were labeled as “polymorphic.” Read depth and read clustering were assessed through custom Python scripts (https://github.com/njdbickhart/python_toolchain/blob/master/metagenomics/plotMagPhaseOutput.py) and IGV^48^ plots.

### Gene cluster prediction and functional annotation

The four assembled metagenomes in FASTA format were used as input for antiSMASH version 5^35^, which predicted the genes using Prodigal^49^. The generated output was used to group BGCs into six different BGC classes: RiPP, NRPS, Terpene, PKS, Saccharide and Others. Also, from the annotated Genbank files, BGCs could be classified into either the “Partial” category (when they were found on a contig edge) or into the “Complete” category (when this was not the case). Finally, predicted BGCs with fewer than 50% of the genes having hits to the best KnownClusterBlast hit, which is obtained from searching all BGCs in the MiBIG database v 2.0^50^ were considered “Novel”. When this condition was not satisfied, the BGCs were classified into the “Known” group.

### Virus and plasmid association analysis

Viral contigs were identified from Blobtools taxonomic assignment for use in the association analysis. Genome completeness of these viral contigs was estimated by the CheckV 1.0 ‘end_to_end’ workflow^40^ (See supplementary table 7). Given the potential novelty of assembled viral genomes in this dataset, the “Not-Determined” and “Medium” completeness viral contigs were not filtered prior to the association analysis. CCS read overlaps and Hi-C link data were used to identify potential host-viral associations as previously described^23^. Briefly, read overlap data consisted of CCS reads that partially mapped to both viral and non-viral contigs. Associative Hi-C links consisted of cases where the number of inter-contig Hi-C pair alignments between viral and non-viral contigs were three standard deviations above the average count for all contigs. Both datasets were compared for overlap, and network plots were generated using the Python NetworkX version 2.5 module. The analysis workflow and network plotting were automated using the following script: https://github.com/njdbickhart/RumenLongReadASM/blob/master/viralAssociationPipeline.py

Plasmids were identified using the SCAPP workflow with the metaFlye HiFi assembly graph (“gfa” file) and aligned short-read BAM files to the final, polished assembly fasta file^36^. The default settings were used apart from the setting of the “-k/--max_kmer” value to “0” in order to disable kmer-based tokenization of sequence reads. SCAPP plasmid nodes were filtered if they were shorter than 5 kb or longer than 1 megabase in length prior to alignment. Plasmid node orthologs in each main assembly were identified through minimap2^43^ alignments and were removed prior to alignment. Hi-C reads were aligned to this modified reference using bwa MEM^42^ and alignment files were converted to BAM format using samtools^47^. The alignment file was used in the aforementioned viral-association workflow script to identify substantial links between candidate plasmids and host contigs. Contig level annotation via the Blobtools^45^ taxify pipeline was used to classify each candidate host by Kingdom. Networks were visualized using the Python NetworkX version 2.5 module.

## Data Availability

The HiFi Sheep dataset, Hi-C reads and WGS short-reads are available on NCBI Bioproject PRJNA595610 at accession ids SRX7628648, SRX10704191, and SRX7649993, respectively. Whole metagenome assemblies and MAG bins for the pCLR and HiFi datasets are available at the following DOI: https://doi.org/10.5281/zenodo.4729049

## Code Availability

The MAGPhase script and codebase are part of the https://github.com/Magdoll/cDNA_Cupcake github repository. Custom scripts used to analyze MAGs and visualize the data are part of the following github repository: https://github.com/njdbickhart/python_toolchain.

## Acknowledgments

We thank Kelsey McClure, Kristen Kuhn, Bob Lee, Jacky Carnahan, and Will Thompson for technical support. DMB was supported by appropriated USDA CRIS project 5090-31000-026-00-D. TPLS and SBS were supported by appropriated USDA CRIS Project 3040-31000-100-00D. We thank Paul J. Weimer for helpful comments and suggestions on the manuscript.

The USDA does not endorse any products or services. Mentioning of trade names is for information purposes only. The USDA is an equal opportunity employer.

## Author Contributions

TPLS and DMB conceived the project with extensive modifications introduced on the advice of IL and PAP. SBS and TPLS were responsible for collecting the sample and generating the sequence data. DB and MK produced the assemblies and conducted a large proportion of reported analysis. VPA and MHM identified biosynthetic gene clusters in the dataset. DMB, AZ and IM identified mobile genetic elements in the sample. ET developed the MagPhase algorithm with inputs from DMB. DMB, TPLS, MK and PAP wrote the manuscript. All authors read and contributed to the final manuscript.

