## Supplementary Figures and Tables for "Generation of lineage-resolved complete metagenome-assembled genomes by precision phasing"

### Precision phasing of lineages within metagenome assemblies using low error long reads

✢ co-corresponding authors

Supplementary Table 1 – Additional file: hifi_total_mag_taxonomy.xlsx

Supplementary Table 2 – Additional file: hifi_phase_bin_association_table.xlsx

Supplementary Table 3: MASH Distance of CLR Clostridia Bins

| | **HIFI 451** | **HIFI 452** | **HIFI 471** | **CLR1 451** | **CLR2 327** | **CLR3 367** |
|---|---|---|---|---|---|---|
| **HIFI 451** | 0 | 0.061 | 0.072 | 0.013 | 0.014 | 0.015 |
| **HIFI 452** | 0.061 | 0 | 0.049 | 0.046 | 0.037 | 0.037 |
| **HIFI 471** | 0.072 | 0.049 | 0 | 0.06 | 0.055 | 0.061 |

Pairwise MASH distances were calculated in a pairwise fashion using the “mash dist” command on kmer sketches derived from each individual bin. All distances were estimated to have p-values less than 0.05 by the program.

Supplementary Table 4: CLR Clostridia Consolidated Bin Statistics

| Assembly | Bin | Ctg count | Total Len | AvgGC | Avg Cov | Stdev Cov | Comp | Contam |
|---|---|---|---|---|---|---|---|---|
| CLR1 | bin3c.451 | 27 | 1973729 | 0.267026 | 43.30088 | 10.40555 | 96.08 | 7.84 |
| CLR2 | bin3c.327 | 26 | 2420085 | 0.261915 | 39.18727 | 8.50268 | 96.08 | 7.89 |
| CLR3 | bin3c.367 | 30 | 2238561 | 0.263730 | 44.01103 | 10.620809 | 96.08 | 7.89 |
| HIFI | bin3c.451 | 7 | 2082786 | 0.267743 | 33.79046 | 12.3909 | 96.08 | 7.89 |
| HIFI | bin3c.452 | 4 | 2077008 | 0.2657 | 21.84393 | 1.526658 | 96.08 | 3.92 |
| HIFI | bin3c.471 | 5 | 2014522 | 0.26316 | 11.06052 | 1.634593 | 96.08 | 5.88 |

Comparison statistics of three resolved HiFi MAGs to their corresponding collapsed bins in the CLR assemblies. The “Ctg count” indicates the number of contigs that comprise each bin, whereas “Total Len” is the cumulative length in bases of the entire bin. The “AvgGC” and “Avg Cov” columns indicate the average GC% and X coverage of short-reads aligned to contigs within each bin. The “Stdev Cov” value is one standard deviation of coverage of short-read coverage in all contigs. “Comp” and “Contam” indicate the percentage of completeness and contamination of each bin using single copy gene metrics, respectively.

Supplementary Table 5 – Additional file: mash_distance_lineage_resolved.xlsx

Supplementary Table 6 – Additional file: clr_dataset_magphase_strain_estimates.xlsx

Supplementary Table 7 – Additional file: viral_contig_checkv_summaries.xlsx

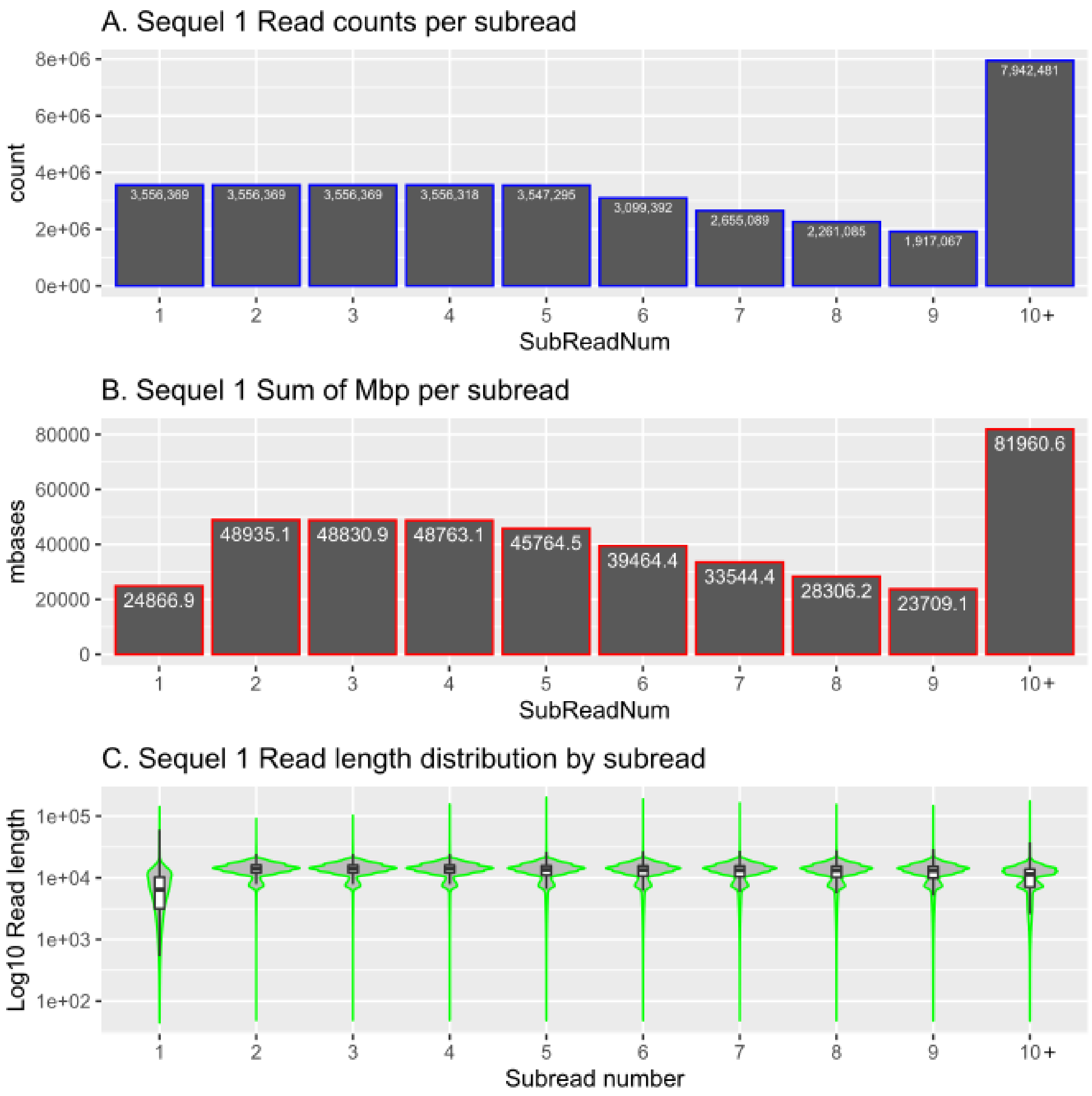


Supplementary Figure 1

PacBio subread counts and statistics for the Sequel I dataset. The count of reads (A), cumulative length (B) and read length distributions (C) of the Sequel I PacBio HiFi reads. Each subread was separated from the original PacBio subread bam files and base qualities were estimated using custom scripts. The X axis represents the sequential count of subreads for each Zero Mode Waveguide, with subread counts at 10 or above being consolidated into one entry (10+).

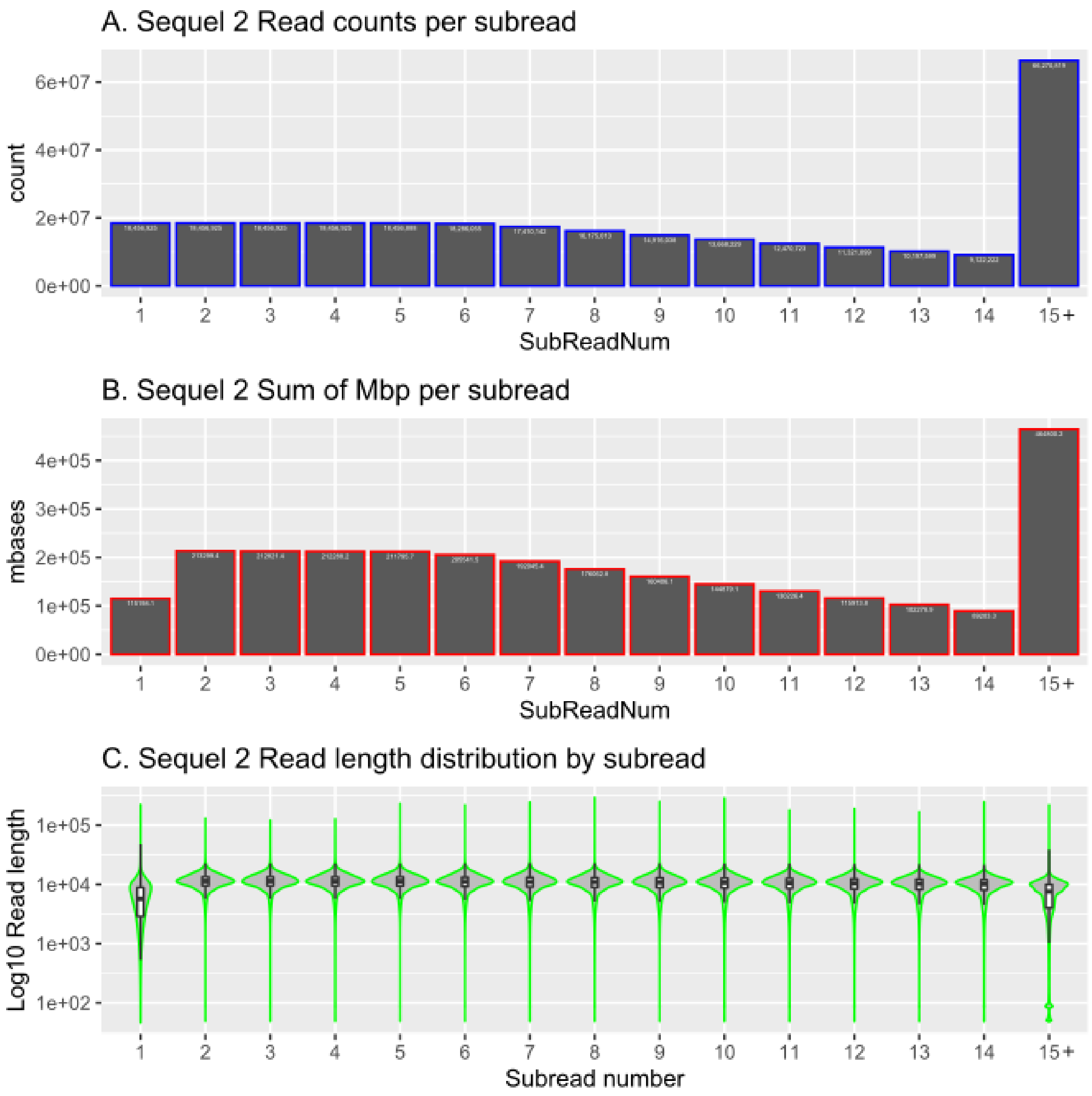


Supplementary Figure 2

PacBio subread counts and statistics for the Sequel II dataset. The count of reads (A), cumulative length (B) and read length distributions (C) of the Sequel II PacBio HiFi reads. Each subread was separated from the original PacBio subread bam files and base qualities were estimated using custom scripts. The X axis represents the sequential count of subreads for each Zero Mode Waveguide, with subread counts at 15 or above being consolidated into one entry (15+).

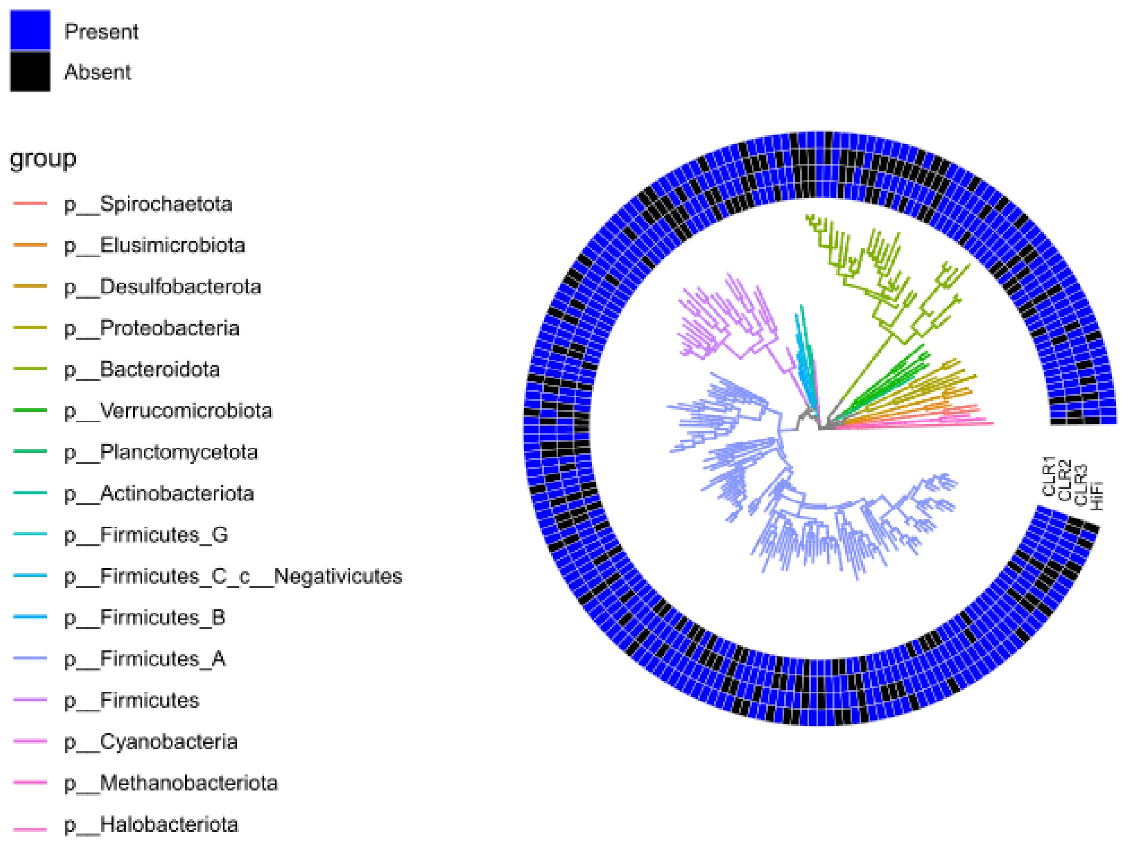


Supplementary Figure 3

A circular dendrogram showing the presence (blue) and absence (black) of GTDB-TK assigned taxonomy to Assembly bins for the HiFi (outermost ring) and CLR (innermost rings, descending) assemblies. Branch nodes were consolidated to Genus-level affiliations when possible. Branch colors were assigned based on Phylum-level classification, with the exception of the Firmicutes, which was sub-divided into separate classes due to its increased diversity relative to other Phyla.

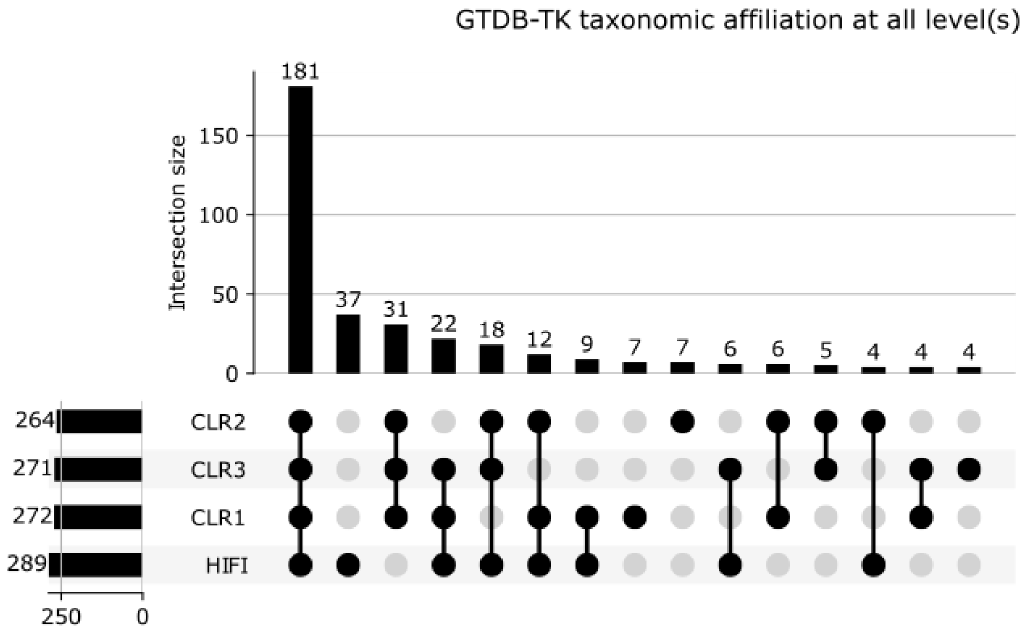


Supplementary Figure 4

An upset plot showing intersections of GTDB-TK taxonomic affiliations for the HiFi and CLR assembly bins. This plot shows the count of intersections of the lowest possible taxonomic classification for each assembly.

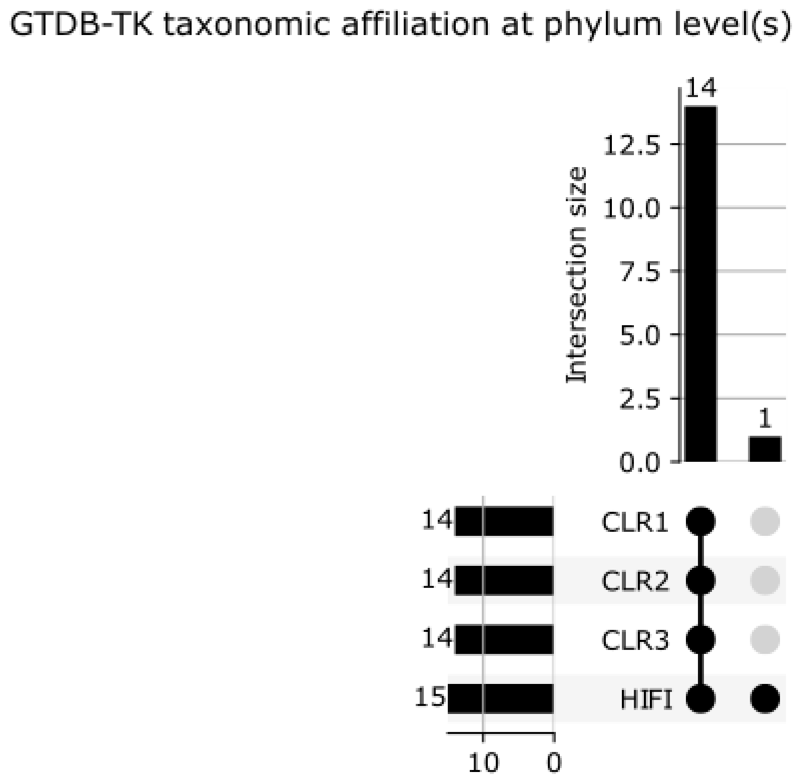


Supplementary Figure 5

An upset plot showing intersections of GTDB-TK taxonomic affiliations for the HiFi and CLR assembly bins. This plot shows the count of unique intersections of Phylum-level classifications only.

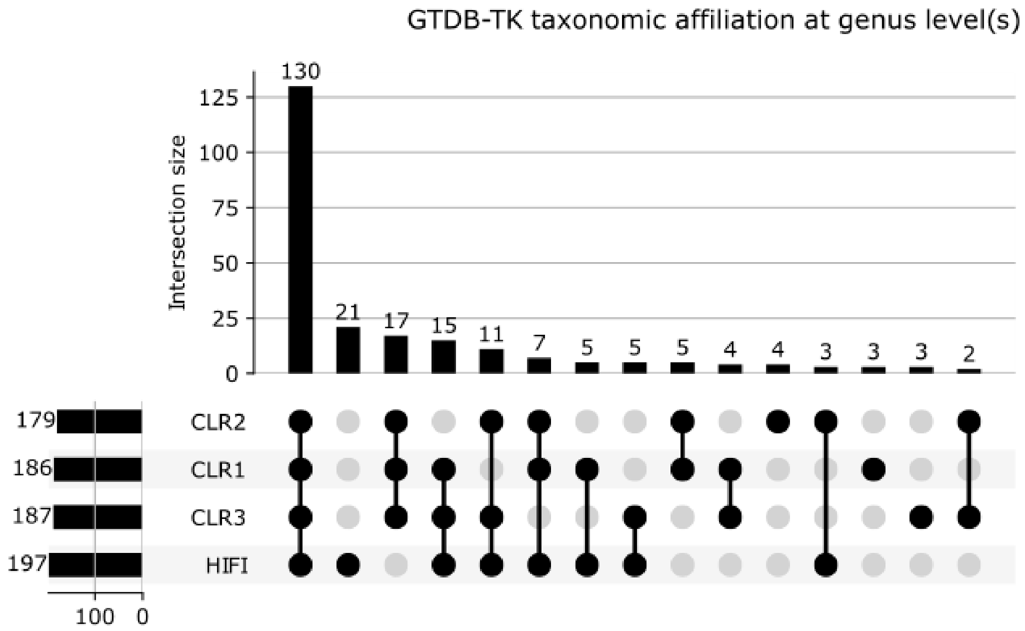


Supplementary Figure 6

An upset plot showing intersections of GTDB-TK taxonomic affiliations for the HiFi and CLR assembly bins. This plot shows the count of unique intersections of Genus-level classifications only.

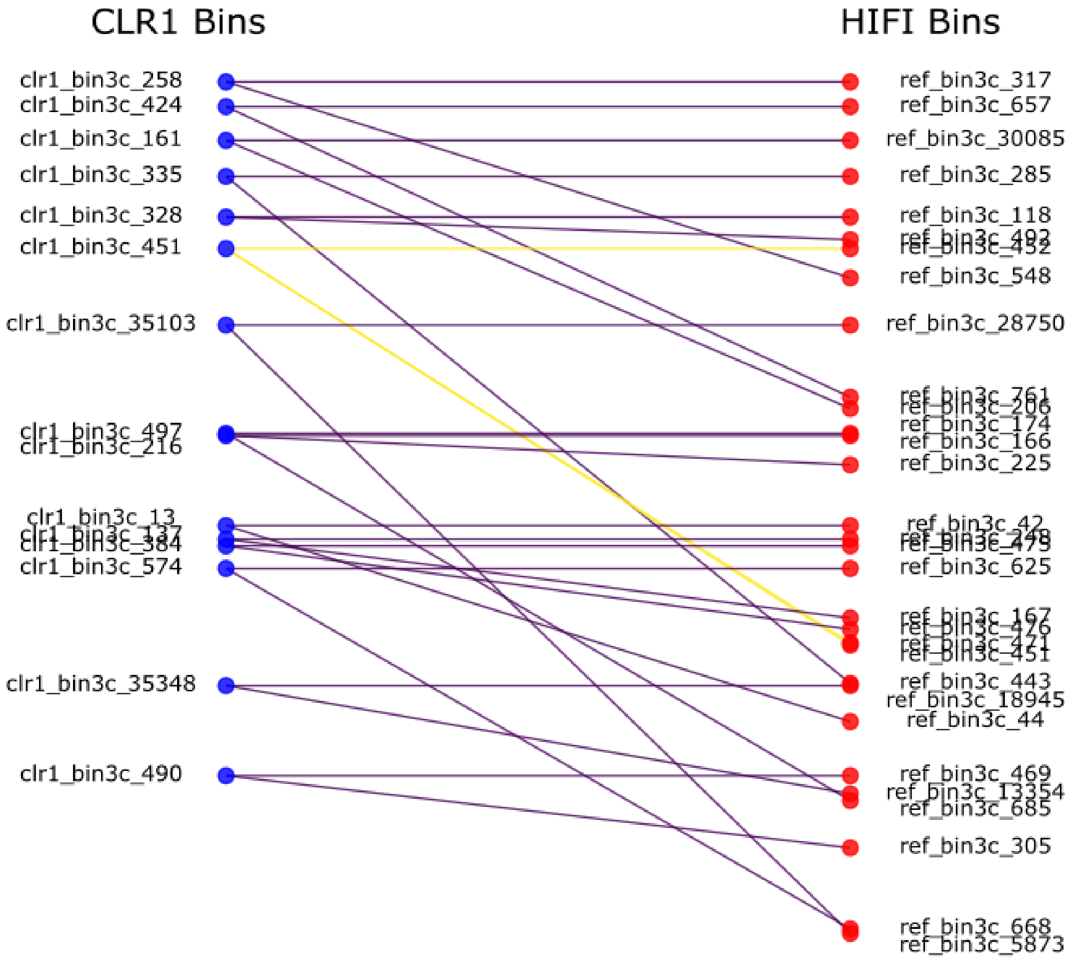


Supplementary Figure 7

A bipartite graph showing the association of predicted collapsed bins in the CLR1 assembly (left) compared to their resolved bins in the HiFi (right) assembly. Collapsed bins in the CLR assemblies were identified as having two or more orthologous bins in the HiFi assembly that had the same GTDB-TK assigned taxonomic affiliation and had MASH distance scores of less than 0.1. Edges in the graph are colored based on the number of associations found for the CLR1 node, with purple indicating two and yellow indicating three associated HiFi bins, respectively.

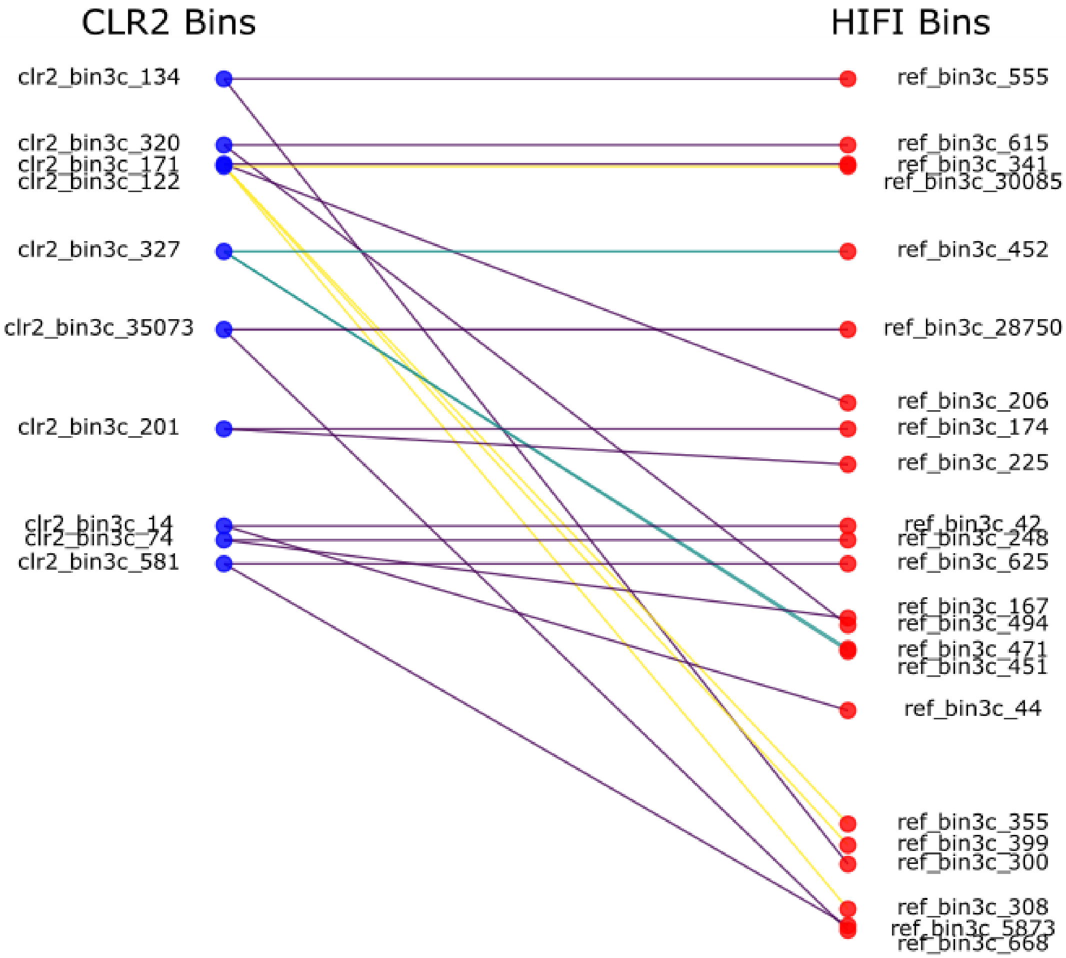


Supplementary Figure 8

A bipartite graph showing the association of predicted collapsed bins in the CLR2 assembly (left) compared to their resolved bins in the HiFi (right) assembly. Collapsed bins in the CLR assemblies were identified as having two or more orthologous bins in the HiFi assembly that had the same GTDB-TK assigned taxonomic affiliation and had MASH distance scores of less than 0.1. Edges in the graph are colored based on the number of associations found for the CLR2 node, with purple indicating two, teal indicating three and yellow indicating four associated HiFi bins, respectively.

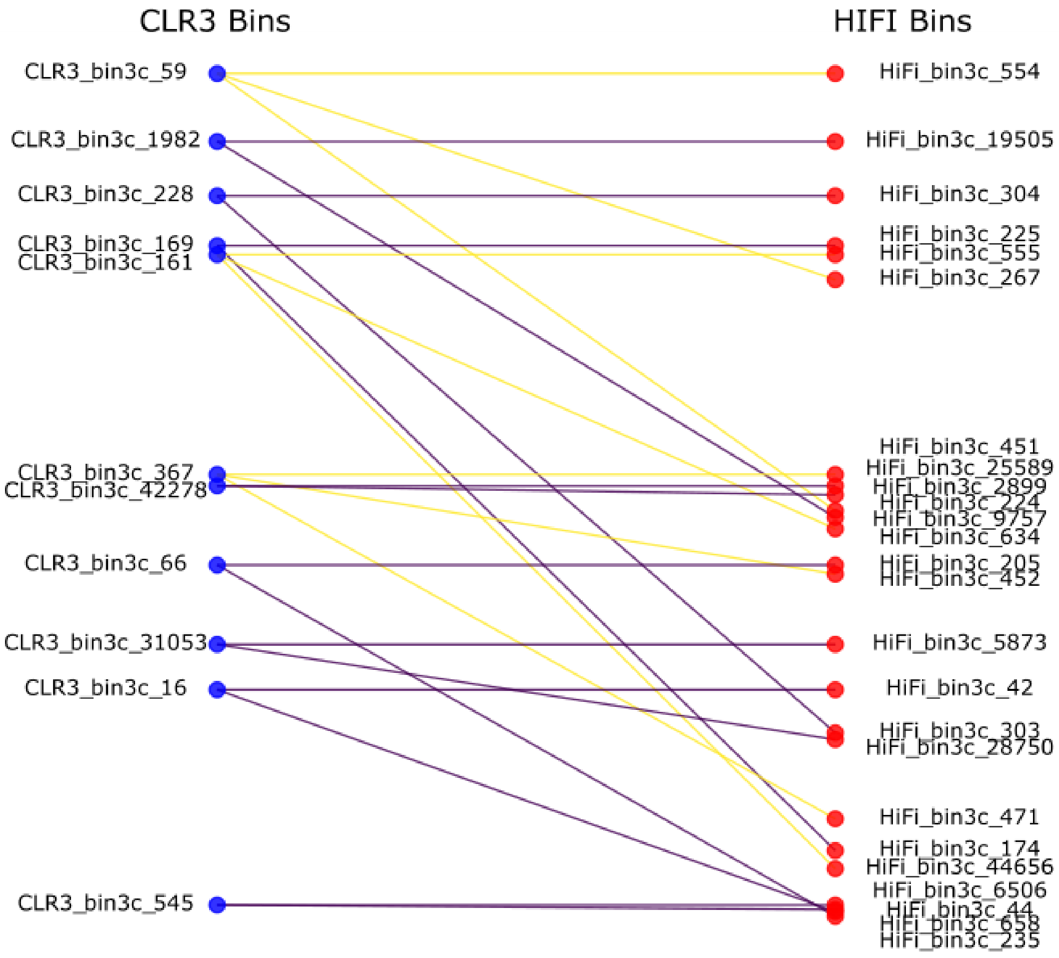


Supplementary Figure 9

A bipartite graph showing the association of predicted collapsed bins in the CLR3 assembly (left) compared to their resolved bins in the HiFi (right) assembly. Collapsed bins in the CLR assemblies were identified as having two or more orthologous bins in the HiFi assembly that had the same GTDB-TK assigned taxonomic affiliation and had MASH distance scores of less than 0.1. Edges in the graph are colored based on the number of associations found for the CLR3 node, with purple indicating two and yellow indicating three associated HiFi bins, respectively.

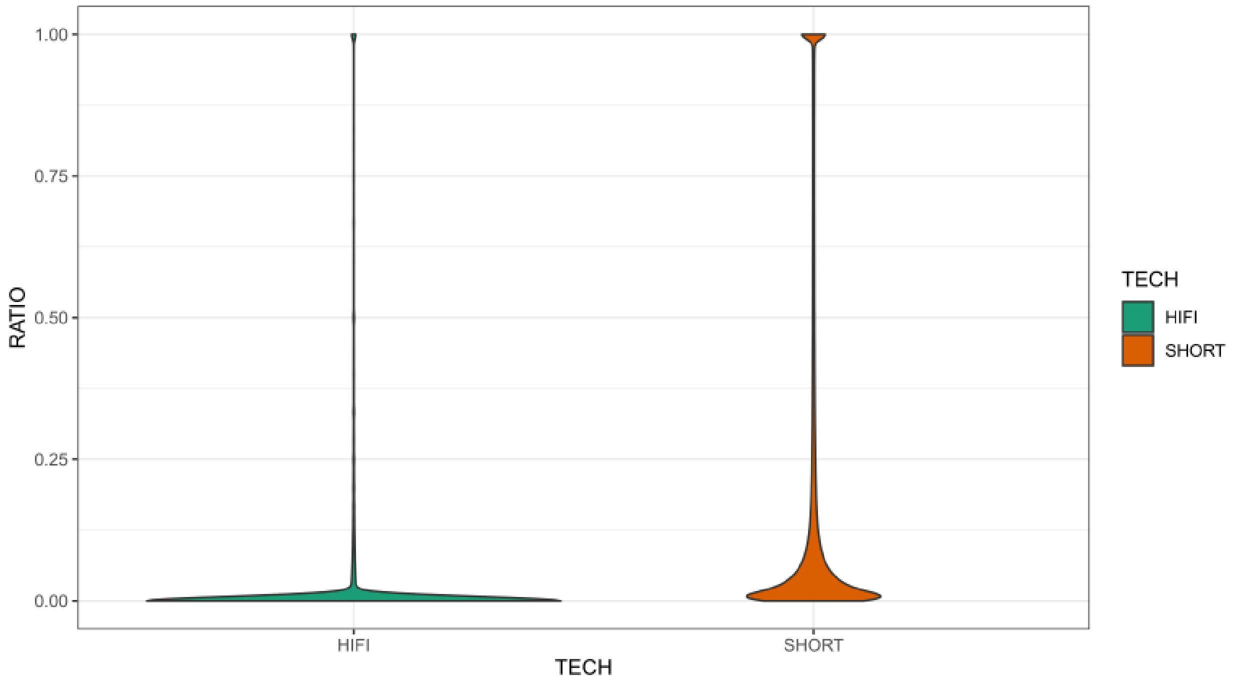


Supplementary Figure 10

The distribution of Map quality score zero read alignments in 5kb windows using HiFi (green) and short Illumina paired-end reads (orange) in the HiFi assembly. Briefly, the entire HiFi assembly was divided into non-overlapping 5kb windows and read alignments were counted from HiFi reads and short-reads in each window. The ratio estimate (y-Axis) represents the proportion of reads that had a Map quality of zero out of the total number of reads within the window. Windows with zero total mapped reads were removed prior to plotting.

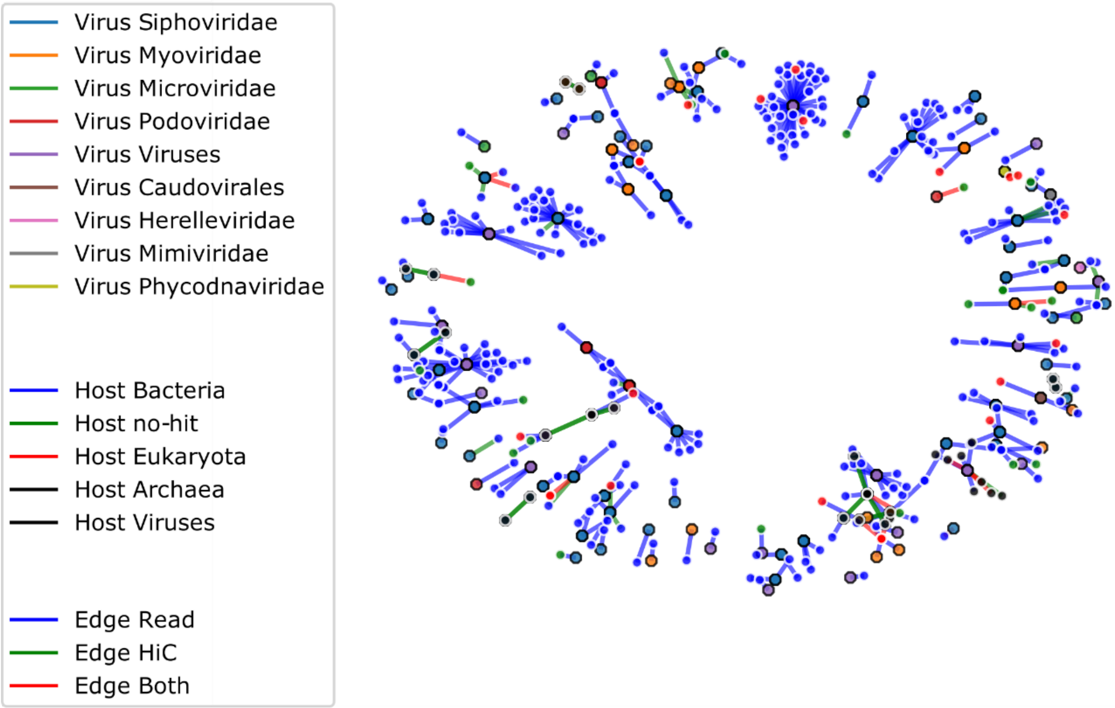


Supplementary Figure 11

CLR1 viral association network plot. Viral contigs identified from Blobtools-assigned taxonomy estimates are represented as hexagonal nodes with black borders, whereas non-viral host contigs are represented as circular nodes with white borders. Edges represent associations identified for each connection, with colors representing the identification of partial HiFi read overlap (blue), Hi-C read links (green) or both types of data (red), respectively.

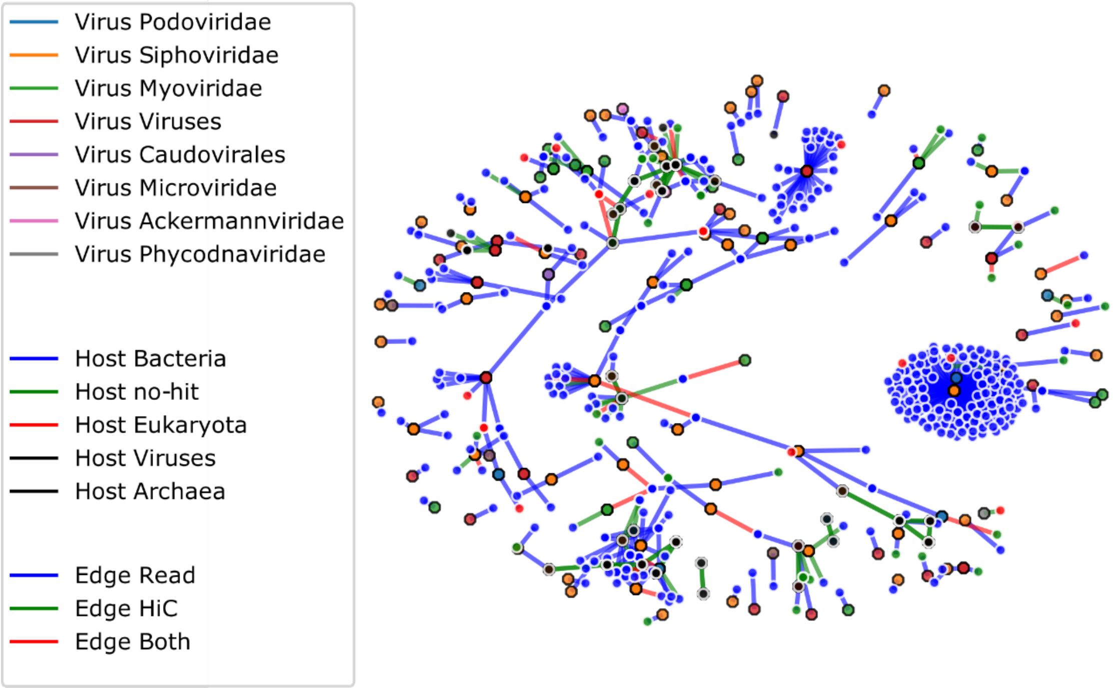


Supplementary Figure 12

CLR2 Viral association network plot. Viral contigs identified from Blobtools-assigned taxonomy estimates are represented as hexagonal nodes with black borders, whereas non-viral host contigs are represented as circular nodes with white borders. Edges represent associations identified for each connection, with colors representing the identification of partial HiFi read overlap (blue), Hi-C read links (green) or both types of data (red), respectively.

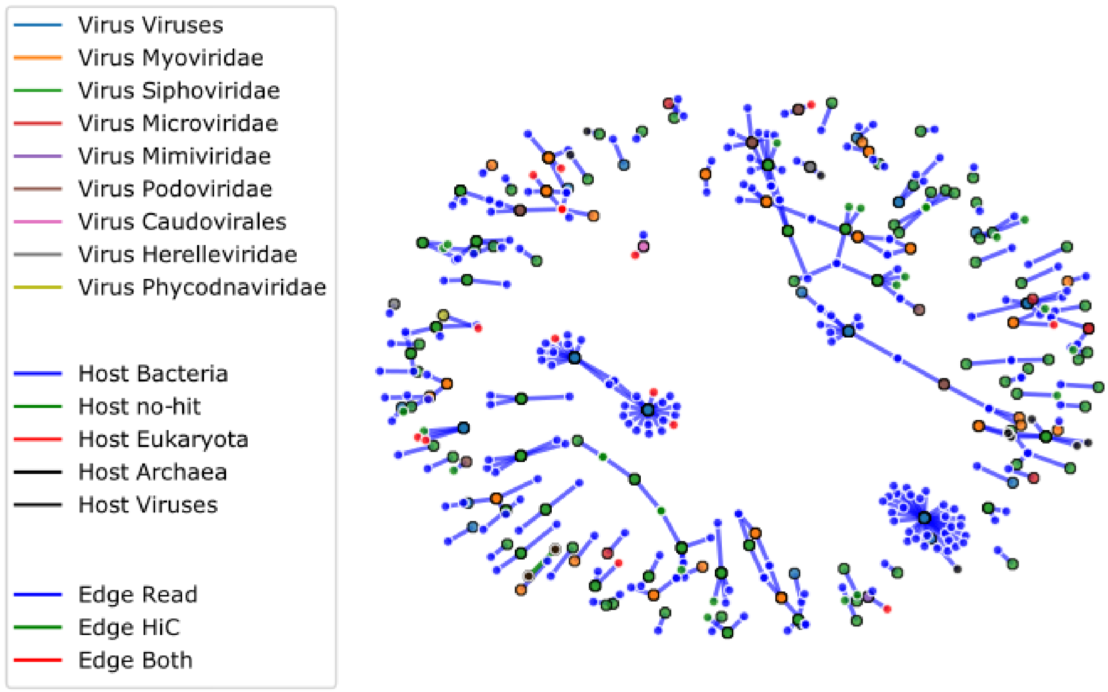


Supplementary Figure 13

CLR3 Viral association network plot. Viral contigs identified from Blobtools-assigned taxonomy estimates are represented as hexagonal nodes with black borders, whereas non-viral host contigs are represented as circular nodes with white borders. Edges represent associations identified for each connection, with colors representing the identification of partial HiFi read overlap (blue), Hi-C read links (green) or both types of data (red), respectively.

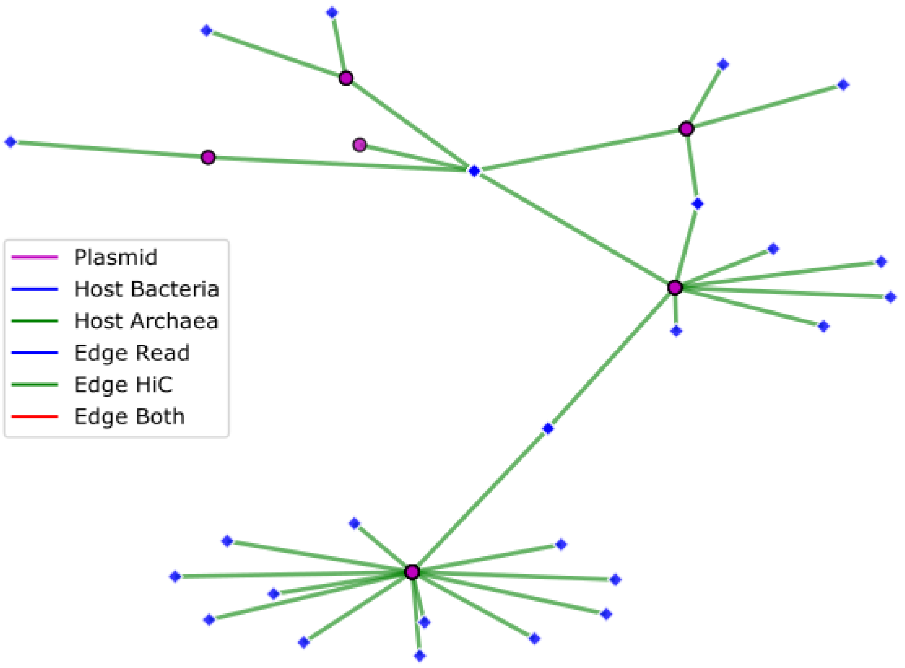


Supplementary Figure 14

The largest subgraph of plasmid-host associations from the HiFi assembly. Plasmid nodes are represented by Hexagonal nodes with black borders and a fill color of purple. Candidate host contigs are represented by diamond nodes with white borders.
